# Behavioral and neural sound classification: Insights from natural and synthetic sounds

**DOI:** 10.1101/2025.01.06.631484

**Authors:** Mattson Ogg

## Abstract

Throughout the course of a day, human listeners encounter an immense variety of sounds in their environment. These are quickly transformed into mental representations of objects and events that guide subsequent cognition and behavior in the world. Previous studies using behavioral and temporally resolved neuroimaging methods have demonstrated the importance of certain acoustic qualities for distinguishing among different classes of sounds during the early time period following sound onset (noisiness, spectral envelope, spectrotemporal change over time, and change in fundamental frequency over time). However, this evidence is largely based on correlational studies of natural sounds. Thus, two additional behavioral (Experiment 1) and EEG (Experiment 2) studies further tested these results using a set of synthesized stimuli (interspersed among a new set of natural sounds) that explicitly manipulated previously identified acoustic dimensions. In addition to finding similar correlational results as previous work (using new natural stimuli and tasks), classification results for the synthesized acoustic feature manipulations reinforced the importance of aperiodicity, spectral envelope, spectral variability and fundamental frequency change for representations of superordinate sound-categories. Analyses of the synthesized stimuli suggest that aperiodicity is a particularly robust cue in distinguishing some categories and that speech is difficult to characterize within this framework (i.e., using these acoustic dimensions and synthetic feature manipulations). These results provide a deeper understanding of the neural and perceptual dynamics that support the recognition of behaviorally important categories of sound in the time windows soon after sound onset.

## I. INTRODUCTION

The environment relays a wealth of physical information that our sensory, neural and cognitive systems distill into representations of objects and events (Bizley & Cohen, 2013; Griffiths & Warren, 2004). This can be viewed as a data reduction stage that is critical for survival and supports our fluent navigation of the external world. The success of object and event recognition is, to a large degree, underpinned by how quickly and automatically recognition occurs. This is especially impressive in audition, where incoming acoustic stimuli require time to develop (Theunissen & Elie, 2014). Thus, early time windows are critical very but limited in the information that can be conveyed shortly after sound onset (Rosen, 1992). A clearer understanding of the rapid processes that influence this perceptual feat will help inform cognitive and neural models of sound recognition as well as downstream speech perception, music perception and auditory scene analysis (see Ogg & Slevc, 2019b for a review). Progress in these areas may in turn inspire improved hearing assistance technologies and machine perception algorithms.

Research in the visual domain provides an example of how useful understanding such temporal dynamics can be. The processing of object related information can be observed in behavioral and neural responses within hundreds of milliseconds following visual stimulus onset (Carlson et al., 2013; Cichy et al., 2014; Thorpe et al., 1996), or given just tens of milliseconds of exposure (Keysers et al., 2001). This has been taken as evidence that feed-forward models can account for the bulk of early visual object processing (Dicarlo et al., 2012). Furthermore, these temporal dynamics provided constraints for neural (Cichy et al., 2014), cognitive (Treisman, 1998), and automated (Serre et al., 2007; Yamins & Dicarlo, 2016; Cichy et al., 2016) models of visual object recognition.

Similar insights might guide models of auditory object and event recognition. Neuroimaging studies have revealed sound-class specific responses in superior temporal cortex (Leaver & Rauschecker, 2010, Lowe et al., 2020; Norman-Haignere et al., 2015; Ogg et al., 2019) that are likely involved in the recognition of speech and musical sounds, as well as materials and actions (Hjortkjær et al., 2017; Lemaitre et al., 2017). However, in addition to encoding specialized, high-level category representations, these studies also identified many cortical regions that appear to respond primarily to relatively low-level acoustic qualities of the incoming stimuli (Kell et al., 2018). An improved understanding of the temporal dynamics of this ostensibly feed-forward acoustic processing and sound recognition stream can shed light on the function of these category-selective responses and better illuminate how human auditory cortex facilitates impressive speech, music and auditory scene perception capabilities.

Beyond classification, responses to the acoustic diversity among natural sounds can illuminate specializations the human auditory system might have evolved in order to support speech and music perception (Smith & Lewicki, 2006). Indeed, results are often described in terms of a delineation between “high-level” category-specific (and ostensibly acoustic-agnostic) processing, and relatively “low-level” processing of acoustic features. Instead, however, recognition (particularly in the context of its rapid temporal unfolding) might be better viewed as a process of transforming pertinent features from incoming stimuli to render a representation (e.g., a manifold; DiCarlo & Cox, 2007) of an object or category (McAdams, 1993; Tsunada & Cohen, 2014; Theunissen & Elie, 2014).

When behavioral and neural responses are studied as a function of a sound’s acoustic qualities, it appears that noisiness, spectral envelope, spectrotemporal variability, and change in fundamental frequency over time all help the auditory system distinguish speech, musical instrument, and human environmental sounds (Gygi et al., 2007; Huang & Elhilali, 2017; Ogg et al., 2017; Ogg & Slevc, 2019a). However, a qualification of these results is that they come primarily from correlational analyses of natural sounds. That is, they involve a collection of neural or behavioral responses to a set of natural stimuli selected to be ostensibly representative of the underlying feature space. Responses are then analyzed as a function of the natural acoustic variability that exists among the stimuli. This yields a view into the features that are particularly important in guiding neural responses and behavior. However, because this approach generally does not involve direct manipulations within the feature space (in favor of indexing the most natural or representative degree of acoustic variability possible), there is a limit to the strength of the inferences that can be drawn about how different features influence the recognition of objects and events. For example, this approach: 1) makes it difficult to isolate the influence of individual acoustic features because it is nearly impossible to orthogonalize each feature within a natural stimulus set, which also, 2) complicates more causal inferences regarding what features a listener relies on when identifying sounds; and 3) may not index the full or relevant range of each feature dimension due to range restrictions that might arise in a given stimulus set (see Thoret, Caramiaux, Depalle & McAdams, 2020 for a related discussion regarding stimulus set limitations).

The studies presented in this report move toward addressing some of these shortcomings among correlational techniques. To do so, two experiments expand on previous sound classification work by introducing explicit manipulations of acoustic features via additional synthesized stimuli. Experiment 1 expands on a previous study that used a behavioral gating paradigm (Ogg et al., 2017) while Experiment 2 follows up on work using temporally resolved neural decoding in M/EEG (Ogg et al., 2020). This allows for a more direct assessment of how influential each of the acoustic qualities previously identified by correlational approaches (noisiness, spectral envelope, spectrotemporal variability, and fundamental frequency variability) are to the early classification and recognition of behaviorally relevant super-ordinate classes (here: speech, musical instrument, and human-environmental sounds).

## II. EXPERIMENT 1: BEHAVIORAL GATING STUDY

One method for examining the temporal dynamics of sound recognition and the information extraction that occurs early in perception is to constrain sound duration and assess subsequent changes to a listener’s discrimination or classification performance. This task is known as a gating paradigm and it has been used to glean insights into the recognition of speech (Grosjean, 1980; Robinson and Patterson, 1995b; Gray, 1942), musical instrument (Robinson and Patterson, 1995a) and everyday action sounds (Lemaitre & Heller, 2013). This task has also been used to study how listeners distinguish among superordinate classes of behaviorally relevant sounds such as speech, music, and materials or tools from an typical human environment (i.e., human-environmental sounds; Bigand et al., 2011; Ogg et al., 2017; Suied et al., 2014). These findings all suggest that the limited information within just tens of milliseconds of sound duration can support above-chance recognition of individual sounds and classification by their superordinate sound classes. Some studies have also reported especially accurate speech recognition under limited duration constraints relative to other classes of sounds (Suied et al., 2014; see also Agus et al., 2012; Isnard et al. 2019; and Moskowitz et al., 2020). Other studies suggest that this effect appears most prominent for vowels, since consonants are often confused with non-speech sounds (e.g., noisy human-environmental sounds; Ogg et al., 2017; Bigand et al., 2011).

In the present study, participants were presented with individual clips of gated sounds asked to indicate whether they had heard a speech, musical instrument or human environmental sound (three-way classification task). Sound duration was very limited in the first block and increased over subsequent blocks as in prior work (Ogg et al., 2017). In addition to a set of natural sounds, the stimulus set included simple synthesized tokens representing controlled manipulations of specific acoustic features that were hypothesized to elicit classification responses for certain sound classes.

### A. Methods and materials

#### 1. Participants

Ninety-six participants (63 female, age: M = 19.5, SD = 1.9, min = 18, max = 27) were recruited from the Psychology Department participant pool at the University of Maryland. They received course credit in exchange for their participation. Three participants were removed prior to analysis due to technical errors (e.g., program accidentally terminated early), and another 6 were removed due to non-compliance with the study procedures (e.g., near-chance performance across duration gates, suggesting arbitrary responses). All participants self-reported normal hearing. Participant recruitment was carried out irrespective of musical ability. The degree and frequency of musical experience in this participant sample (72% received some degree of musical training: M = 5.8, SD = 3.3) was representative of large undergraduate university populations (Schellenberg, 2006; Corrigall et al., 2013) and similar to previous reports within this community (Ogg et al., 2017; Ogg & Slevc, 2020).

#### 2. Stimuli

##### a. Synthesized stimuli

All stimuli were generated via a digital synthesizer in Ableton Live (“Operator” Live version 9; Ableton, Berlin, Germany) with all settings turned off or set to their default position (single oscillator source with no frequency modulation of that source) except where noted in the description of each manipulation below. Each sound token was initially generated with a 300-millisecond duration. Limited time scales such as these are more suited to relay spectral and harmonic information rather than information from temporal envelopes due to inherent physical trade-offs in spectral and temporal resolution (Moore, 2012). Thus, all synthesized stimuli were generated with steady state temporal envelopes. Each acoustic manipulation was characterized by three values that were spaced two large (approximately logarithmic), steps apart along each acoustic dimension. This spacing was chosen to achieve perceptually salient changes that might similarly distinguish speech, musical instrument and human-environmental sounds.

Whenever a given dimension was manipulated, every effort was made to control changes in the other dimensions to isolate the influence of individual acoustic cues. This was implemented either by eliminating other dimensions where possible or by holding them constant. However, as with naturally occurring sounds, these synthesized stimuli are subject to certain physical constraints that make total separation of some acoustic dimensions difficult to achieve. For example, high noisiness also increases spectrotemporal variability because the signal is uncorrelated with itself from moment to moment. Since the goal of this approach was to generate stimuli to yield controlled representations of acoustic properties of natural sounds, some co-variation among particular acoustic dimensions in the synthesized stimuli was reasonable as long as that co-variation was also present in the natural sounds. Similarly, it was necessary to include some interactions among features to address constraints that are also present in natural sounds (e.g., it is not possible to manipulate a spectral envelope without some carrier signal to act upon).

The aperiodicity manipulation served as the carrier for the other spectral manipulations. All stimuli were based on either a white-noise or sawtooth-wave carrier, the latter having 64 equal amplitude harmonics (fundamental frequency set to E♭_4_ or 311 Hz). The white noise source represented a broadband sound without any harmonic components, while the sawtooth wave had many harmonics, and is frequently used as a source waveform for synthesized speech and instrument sounds^1^.

These different carriers (i.e., the synthesizer’s oscillator) were submitted to modifications addressing the other acoustic dimensions of interest as follows. The spectral centroid manipulation constrained the signal’s spectral bandwidth via a low-pass filter with a low (1000 Hz), medium (4000 Hz) or high (16000 Hz) frequency pass-band (i.e., where the filter cut-off reaches −12dB given a 12dB/octave roll-off) to manipulate the overall spectral envelope (centroid) of the sounds. Tones in the spectral centroid condition had a constant filter pass-band frequency and fundamental frequency to control for the changes in other dimensions (spectral and fundamental frequency variability, respectively).

The spectral variability manipulation introduced a low (+/− 190 Hz), medium (+/− 750 Hz), or high (+/− 3000 Hz) degree of modulation in the pass-band frequency of the low-pass filter. Modulations were implemented via a 150-millisecond linear up-down sweep of the pass-band. This up-down sweep pattern was chosen because it resembles the dynamics of many initial excitatory acoustic patterns in the natural sound set, such as from a brass instrument onset, a liquid sound (e.g., a drip or splashing), or a mouth opening and closing. Each filter sweep was centered around 4000 Hz and the tones maintained a consistent fundamental frequency to control for changes in the other dimensions (median spectral centroid and fundamental frequency variability, respectively).

Finally, the fundamental frequency manipulation introduced a low (25 cents, or a quarter semi-tone), medium (100 cents, or one semi-tone) or high (200 cents, or two semi-tones) degree of change in the fundamental frequency of the sawtooth waves via a linear downward sweep of the oscillator’s pitch control. A downward sweep was chosen because most of the fundamental frequency variability observed in the speech tokens in prior work was in a downward direction (Ogg et al., 2017; Ogg & Slevc, 2019a). Each sweep unfolded over the stimulus’ entire 300 milliseconds duration. All of the tones were low-pass filtered at a constant 4000 Hz frequency to control for changes in the other dimensions (spectral centroid and spectral variability). This manipulation was not crossed with the noise condition because the noise carrier did not possess a fundamental frequency that could be manipulated.

A schematic of this design and a depiction of the stimuli at each step along the manipulated dimensions can be found in Figure 1 along with indications for the super-ordinate sound class each stimulus was hypothesized to represent. Musical instrument and human-environmental sounds were hypothesized to be represented by low and high extremes along these feature dimensions, respectively. Meanwhile, speech tokens were hypothesized to be represented by stimuli that occupied the space between these extremes. Less strong hypotheses exist for the stimuli which might plausibly correspond to multiple categories.

**Figure 1.**
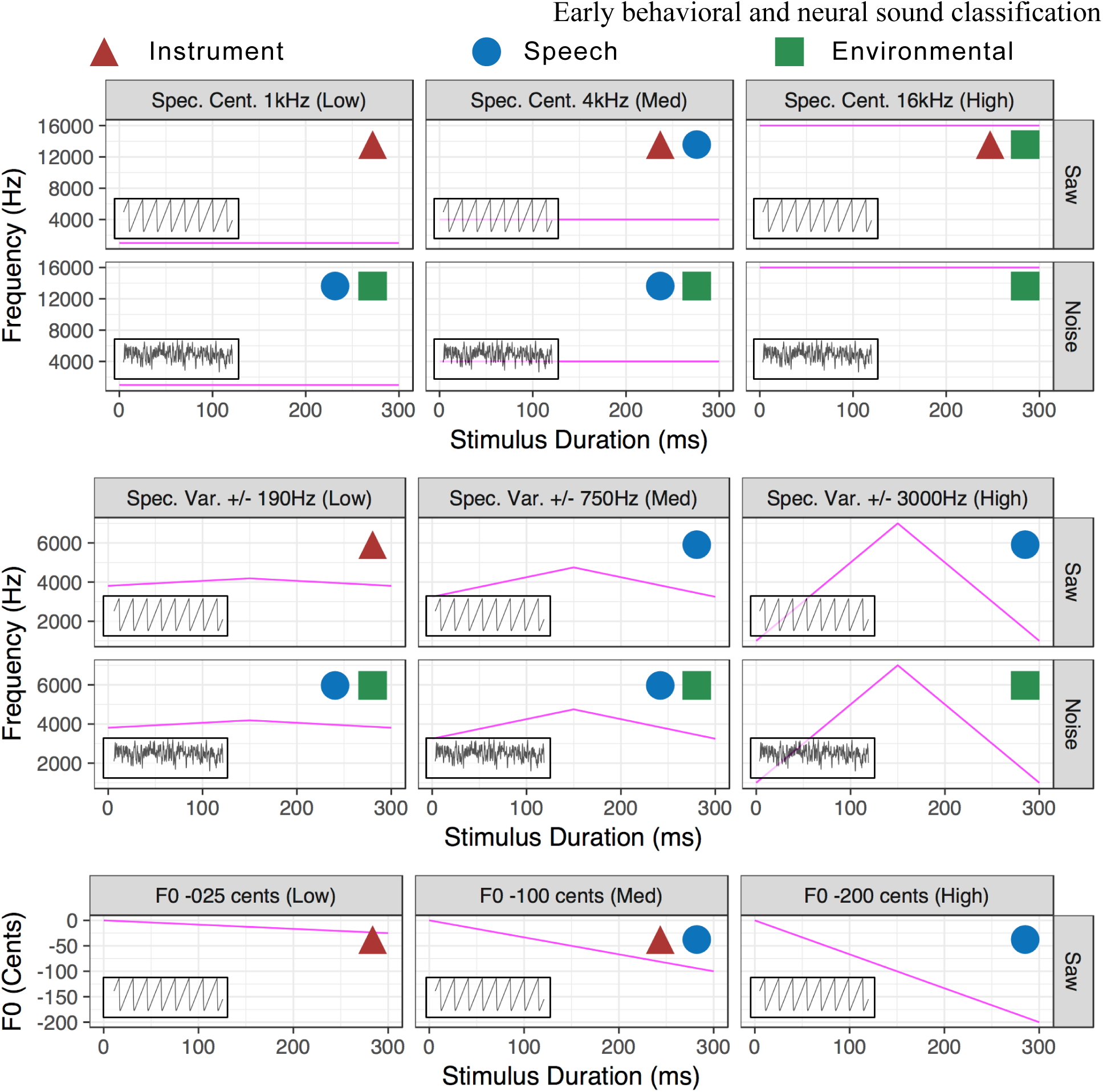
Schematic of synthesized stimulus manipulations. Magenta lines indicate the trajectory of the stimulus’ low-pass filter or fundamental frequency (vis-a-vis their respective y-axes) throughout the course of each 300 ms stimulus. These are plotted from low (saw carrier) to high (noise carrier) aperiodicity (top and bottom rows within each panel, respectively) and low to high manipulation levels (left to right columns within each panel) for spectral centroid, spectral variability, and fundamental frequency variability. Colored shapes in the top right of each panel indicate the sound class each stimulus is hypothesized to represent. Insets are a schematic of the acoustic carrier of each stimulus in a given condition (panel).

##### b. Natural stimuli

The 15 synthesized stimuli described above were added to a pool of 108 speech, musical instrument and human-environmental sound recordings. This was for two reasons. First, the natural stimuli operationalized each sound class for the listener. Second, including these natural sound tokens enabled a built-in test of prior findings by running the same analyses on the trials containing natural sounds. This pseudo-replication serves to validate the manipulations used among the synthesized stimuli.

The natural stimuli were novel sound files similar to (but not the same as) prior related work (Ogg et al., 2017; 2020). This stimulus set was generated from different sound databases, new recordings or new draws of sounds from previous databases. However, the natural stimulus set as a whole was designed to meet the same general criteria as previous studies, which was to sample the acoustic variability within and between these sound classes. The full list of sounds can be found in Figure 2.

**Figure 2.**
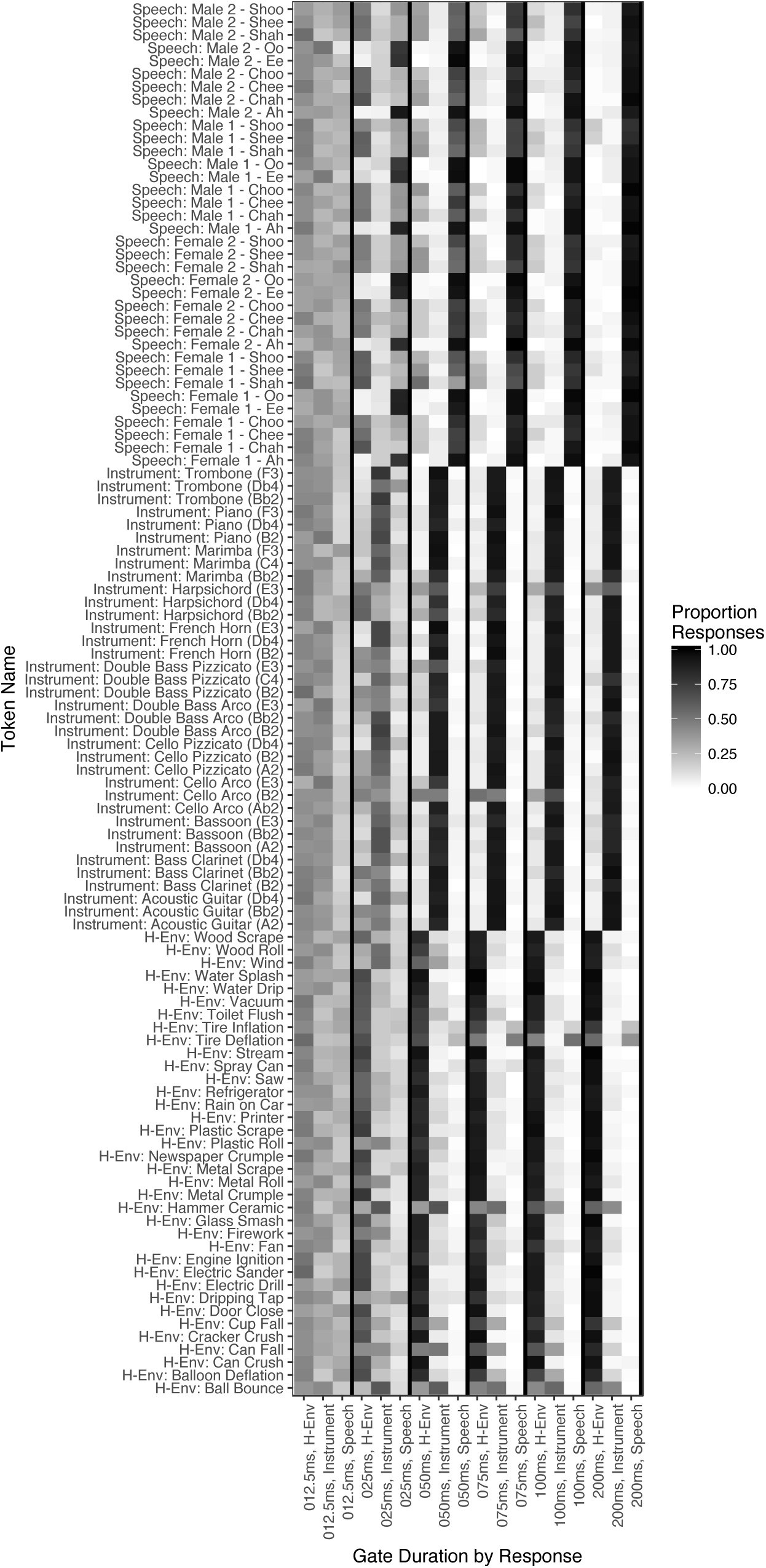
Item-level confusion matrix of responses for natural stimuli by each response class and duration in Experiment 1.

Speech tokens were recorded in-house and comprised the vowels /ɑ/, /i/ and /u/ spoken by themselves or following the consonants /tʃ/ or /ʃ/ (e.g., “chah,” “shah,” etc.). Speech stimuli were spoken by two males and two females (native English speakers, all of whom grew up in the Mid-Atlantic region of the United States). The inclusion of both vowels and consonants in the stimulus set helps form a more complete picture of how listeners treat these diverse parts of speech (cf. Ogg et al., 2017). During recording, speakers read a randomized list of the above consonant-vowel (or vowel) utterances. Speakers were asked to elongate the vowel of each utterance in time and to keep their speaking pitch as steady as possible, but to otherwise speak normally. Recordings were made in a sound treated room with high fidelity equipment (studio quality condenser microphone and analog-to-digital recording interface, at least a 16 bit-depth and 44,100 Hz sampling rate; see Ogg et al., 2017 for further description of the recording equipment). The median fundamental frequency of each utterance was calculated using the YIN algorithm (de Cheveigné & Kawahara, 2002), and the closest musical notes were identified for matching the musical instrument stimuli: A_2_, A♭_2_, B_2_, B♭_2_, C_4_, D♭_4_, E_3_, and F_3_ (104.90 Hz to 282.08 Hz).

Musical instrument tokens were drawn from the McGill University Master Samples database (Opolko and Wapnick, 2006), which contains high quality recordings of a diverse array of instruments playing their full range of notes. Instruments were selected to span the orchestral timbre palette with the constraint that the instruments’ fundamental frequencies could match the male and female speaker vocalizations. Notes from a marimba, trombone, french horn, bassoon, bass clarinet, double bass arco, double bass pizzicato, cello arco, cello pizzicato, piano, harpsichord, and acoustic guitar were used to meet these criteria. Recordings of three notes were obtained for each instrument corresponding to three randomly selected notes from the set of vocalization fundamental frequencies. Note selection for the instruments was constrained such that at least one note corresponded to an utterance from a speaker of each gender (thus approximately an octave apart).

Environmental sound tokens were manipulations or excitations of everyday objects and materials selected from the Carnegie Mellon University Sound Events and Real-World Events Databases (2008; Vettel, 2010) and the BBC Sound Effects Library (1997, BBC Worldwide, London, United Kingdom). Six sounds represented each of the following methods of acoustic excitation (approximating previously outlined taxonomies described by Gaver, 1993 and Lemaitre & Heller, 2013; 2012): air, deformation, impact, mechanical, movement, and liquid sounds. These included sounds such as air exiting a balloon, glass breaking, an engine ignition, or water dripping.

All stimuli (both the natural and synthesized sounds) were edited to begin at their onsets, defined here as 5-milliseconds prior to where the stimulus’ absolute amplitude reached 10% of its maximum. Stimuli were then edited to 12.5, 25, 50, 75, 100 and 200-millisecond durations and 2.5 millisecond cosine onset and offset ramps were applied (a 300-millisecond version was also generated for the familiarization block and for Experiment 2). All the stimuli (natural and synthesized) were then normalized to the same root-mean-square level.

The acoustic features of the stimuli were extracted using similar methods as Ogg et al. 2020 and Ogg & Slevc, 2019a. See Table 1 for a detailed description of each feature along with notes regarding calculation and interpretation. Windowed or temporally varying features were extracted in 5 millisecond increments and summarized using the median or IQR of values across timesteps similar to previous work.

**Table 1.** Description of Acoustic Feature Derivation and Interpretation.

| Feature | Description | Interpretation |
| --- | --- | --- |
| Log-Attack-Time<sup>1</sup> | Log of the time difference between attack onset and ending | Lower values = faster onset time |
| Temporal Centroid<sup>1</sup> | Center of gravity of the energy envelope | Lower values = earlier temporal centroid |
| Spectral Centroid<sup>2</sup> | Center of gravity of the spectral (ERB) envelope | High values = higher frequency centroid |
| Spectral Flatness<sup>2</sup> | Ratio of geometric and arithmetic means of the spectral (ERB) envelope | Measures noise/harmonic content. Higher values are flatter/noisier |
| Spectral Variability<sup>2</sup> | 1 minus the correlation of ERB channel spectra between successive timepoints | Higher values = more variable envelope |
| Aperiodicity<sup>3</sup> | Amount of aperiodic energy in the signal | Higher values = more aperiodic |
| ERB Energy<sup>2,e</sup> | Amount of energy in the spectral representation at each timestep (frame) | Sum of squared amplitudes in the spectral representation. |
| Raw ERB cochleagram<sup>2,e</sup> | Raw ERB cochleagram representation of each of 77 channels: 30 Hz to 16 kHz | Energy in each channel over time. |
| Mod. Power Spectrum<sup>4,e</sup> | 2D-FFT of Gaussian spectrogram | Degree of joint spectral/temporal modulation rates |

Derived based on: 1) Energy envelope, 2) ERB cochleagram representation in the Timbre Toolbox (Peeters et al., 2011; Kazazis et al., 2017), 3) YIN (de Cheveigné & Kawahara, 2002), or 4) modulation power spectrum (Elliot and Theunissen, 2009); e) Used in Experiment 2 only. Adapted from Ogg & Slevc (2019a).

#### 3. Apparatus

Instructions, stimulus presentation and response logging for the experiment were conducted using PsychoPy (version 1.82, Peirce, 2007) on an Apple computer (model number: MC508xx/A; Apple Computer, Cupertino, CA). Participants performed the study individually in a private room. Stimuli were played back via Sennheiser HD 202-ii headphones (Sennheiser Electronics, GmBH, Wedemark, Germany) at approximately 77-dBA.

#### 4. Procedure

Participants were told they would hear a wide variety of sounds and that they would be asked to push a button to indicate whether they heard a speech, musical instrument, or environmental sound. A written description of an example from each sound class was provided. Participants were told that the sounds they would hear might be very short and to make their best guess if they were not sure what kind of sound they heard. After reading the instructions, a passive (no response required) familiarization block began for the 108 natural stimuli which involved playback of each natural sound in the stimulus set (played twice during each familiarization trial: 300-millisecond duration, 1 second inter-onset-interval). During each trial in the familiarization block, the sound’s label (e.g., “Bassoon”) and class (e.g., “Instrument”) was displayed for 2.5 seconds. A one second inter-trial-interval then followed before the next familiarization trial.

After the familiarization block, participants began the gated sound classification task. During each block, all 123 (natural and synthesized) stimuli were presented in a random order and on each trial the participant was asked to indicate what kind of sound they heard via button press. Instructions appeared on-screen (“What kind of sound did you hear? C if speech; B if instrument; M if environment”) 500 milliseconds before sound onset. The participant’s subsequent response triggered a 1 second inter-trial-interval before the next trial started. Each block of trials consisted of the same gate duration, and the order of durations proceeded sequentially across blocks from the 12.5-millisecond gate to the 200-millisecond gate. Participants were given the opportunity to take a short break halfway through. Note that participants were never informed of the synthesized stimuli nor introduced to them prior to the first block of trials.

#### 5. Data analysis

Analysis of the natural stimuli (i.e., limited to the subset of trials that contained natural sounds) was carried out in a similar fashion as in Ogg et al. (2017). This was to validate that the feature manipulations of the synthesized stimuli, which were based on the results of past studies, were still valid for this new dataset. Modifications were made to that analysis owing mainly to the three-choice response measure of the present study (as opposed to the go/no-go task used in the previous study). Trial-level accuracy was analyzed as well as block-level *d′* scores. Within a given block (i.e., each duration gate) *d′* was calculated for each class such that a correct class response was coded as a hit (i.e., a correct class response when that class was present) and an incorrect class response was coded as a false alarm (i.e., a given class response when a different class was present). Counts of hits and false alarms were converted to probabilities by dividing by the number of possible hits and false alarms (36 and 72, respectively; ceiling and floor counts adjusted by 0.5).

Block-level *d′* scores were analyzed via linear mixed-effects regressions (all mixed-effects models fit using ‘lme4’ in R; Bates et al., 2015 with statistical significance assessed where applicable using the ‘lmerTest’ package; Kuznetsova et al., 2017) with predictors for sound class (dummy coded: speech, instrument and human-environmental with speech as the intercept), gate duration (which was log-transformed and z-normalized), musical training (z-normalized musical training subscale of the Gold-MSI; Müllensiefen et al., 2014) and the interactions among these variables. Trial-level accuracy was modeled using a generalized binomial mixed effect regression (with a logit link function) with the same predictors as the model of *d′* scores. In both models, participants were entered as intercepts into the model’s random effects, with gate duration nested as a random slope within participants (the slopes and intercepts were constrained to be uncorrelated with one another). The trial-level accuracy model also contained an intercept for items (all 108 regardless of duration) with gate duration nested as a random slope (also constrained to be uncorrelated with the random intercept of items). Model formulas are indicated below each table of results throughout this report.

For analyses of acoustic features of the natural sounds, the three-choice response variable was recoded via “one-hot” encoding into three new binary outcome variables (one binary variable for each sound class: speech, music or human-environmental). These indicated whether a trial contained a response for a given sound-class (1) or not (0). These were used as outcome variables in three subsequent all-subsets regression analyses (using generalized binomial mixed effects models and the ‘MuMIn’ R package, Bartoń, 2017) to identify the most parsimonious set of acoustic features associated with participants’ responses for each class. These analyses ranked models of all possible combinations of the acoustic features according to their fit to the data (conservative BIC criterion), and the best fitting models (within 3 BIC of the top) were retained and averaged (Burnham and Anderson, 2004; Whittingham et al., 2006).

The all-subsets regression analyses were carried out in two stages^2^. The first stage was restricted to individual (z-normalized) acoustic feature predictors without interactions with duration (i.e., the influence of each feature irrespective of the duration gate/block). In the second stage, another all-subsets regression was carried out that included only predictors for the top-ranked features from the first stage (i.e., those features present in models within 3 BIC of the top-ranked model) as well as their interactions with duration (which was log-transformed and z-normalized). The random effects for these models contained intercepts for participants and items with gate duration nested as a random slope within each intercept, which was constrained to be uncorrelated with the intercepts.

Finally, the synthesized stimuli were analyzed using the subset of trials that contained only those stimuli (i.e., no trials with natural sounds) in a series of generalized binomial mixed effects regressions (with a logit link function). These models were also fit to three “one-hot encoded” binary outcome variables for each class response. Separate models were fit for each of the three manipulation conditions and for each sound class (except for the aperiodicity manipulation which interacted with the other manipulations), for a total of nine models. Each of these models was fit only to the subset of trials relevant to each manipulation. These models contained predictors for duration (log-transformed and z-normalized), the aperiodicity condition (i.e., the acoustic carrier with 2 contrast coded levels: low aperiodicity/sawtooth and high aperiodicity/noise, with the sawtooth condition as the reference level; rows of the sub-panels in Figure 1), and a predictor for one of the other manipulations for spectral centroid, spectral variability or fundamental frequency (each with three levels: low, medium, or high, dummy coded with low level designated as the intercept; columns in Figure 1). Models also included the two and three-way interactions among these variables. Note that models of the fundamental frequency manipulation did not contain the aperiodicity predictor because there was no noise stimulus for that manipulation (all pitch manipulations acted upon the sawtooth wave carrier). The random effects structure of these models contained random intercepts for participants with a slope for duration nested within participants, and these random effect slopes and intercepts were constrained to be uncorrelated. Preliminary analyses did not indicate an appreciable influence of musical training when this variable (and its interaction with the other three predictors) was included in these models, so musical experience was excluded from the report of these models for simplicity^3^.

### B. Results

Classification responses for three sound classes were analyzed for this gating task where sound duration was at first greatly constrained (12.5 milliseconds) then increased over the course of six blocks of trials (200 milliseconds). Responses (classification of a clip as speech, music or human-environmental) were obtained for a set of natural sound stimuli as well as a set of randomly interspersed synthesized stimuli designed to directly manipulate a set of relevant acoustic dimensions. The acoustic manipulations characterized by the synthesized stimuli build off of previous results based on analyses of natural sounds (Ogg et al., 2017), so this section will first briefly revisit those analyses within this new task and stimulus set to validate the applicability of the manipulations for these data.

#### 1. Analyses of the natural sound tokens

Participants’ responses for each stimulus and sound-class across duration gates (blocks of trials) are summarized in confusion matrices in Figures 2 and 3, respectively. Participant accuracy on this three-choice task is summarized in terms of proportion correct (by block for the sounds of each class) in Figure 4 and in terms of *d’* scores (calculated by participant for each block and class response) in Figure 5. In general, these findings are similar to previous results (Ogg et al., 2017), although discriminability (*d’*) for speech sounds was higher than other categories across duration gates, which is perhaps expected given the inclusion of isolated vowel sounds in the stimulus set (compare experiments 1 and 2 in Ogg et al., 2017), which listeners have been shown to be especially accurate at classifying as speech (Agus et al., 2012; Suied et al., 2014).

**Figure 3.**
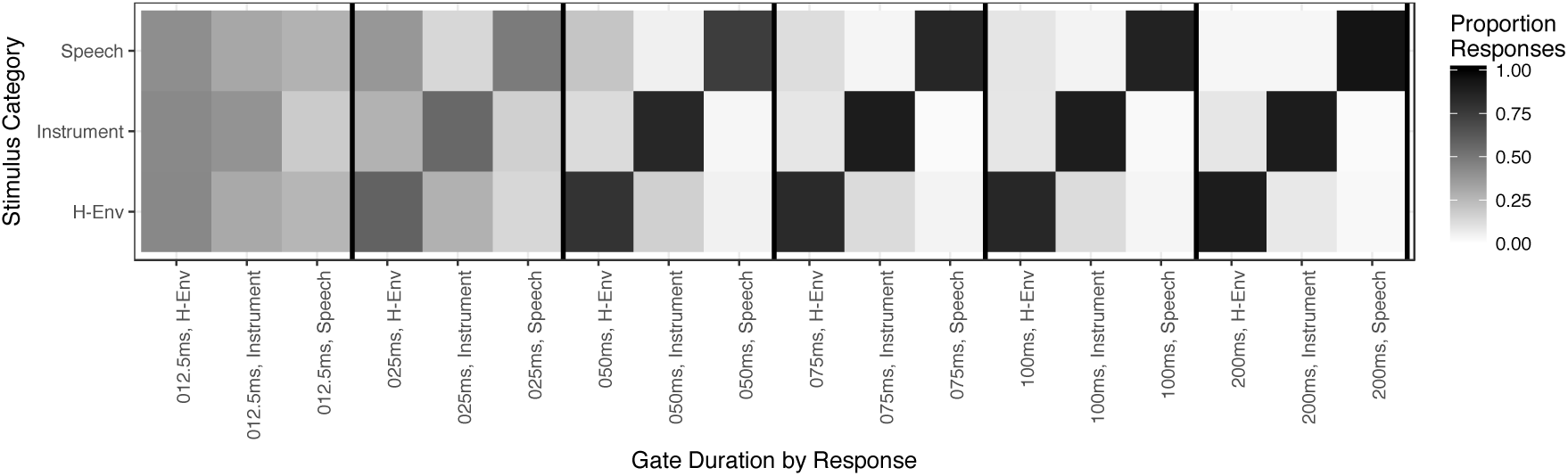
Sound-class-level confusion matrix of responses for the natural stimuli by each response class and duration in Experiment 1.

**Figure 4.**
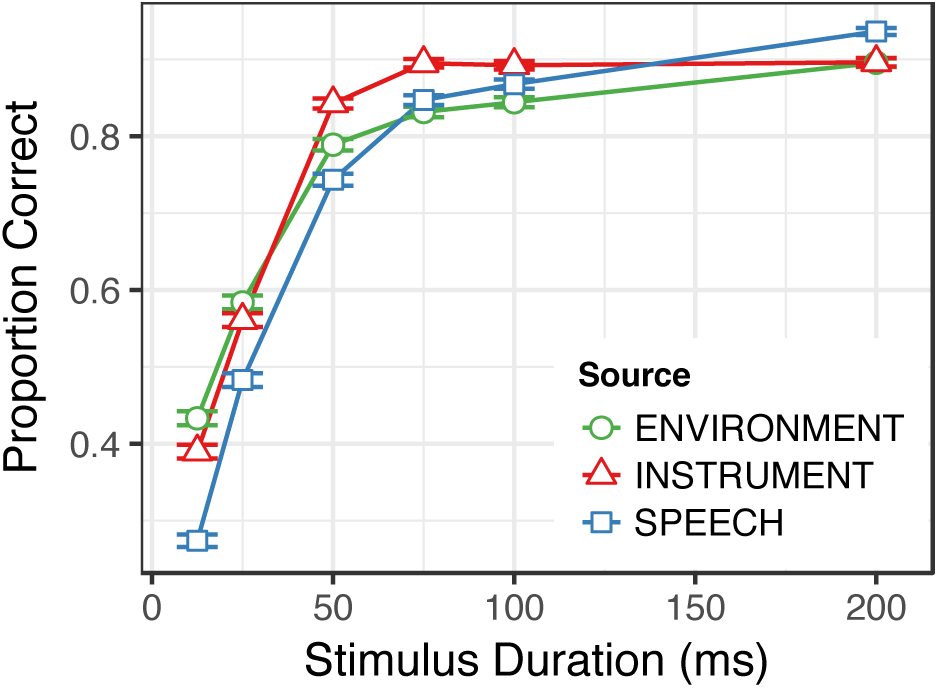
Proportion of correct responses across participants in each block of Experiment 1 displayed by target sound-class as a function of gate duration.

**Figure 5.**
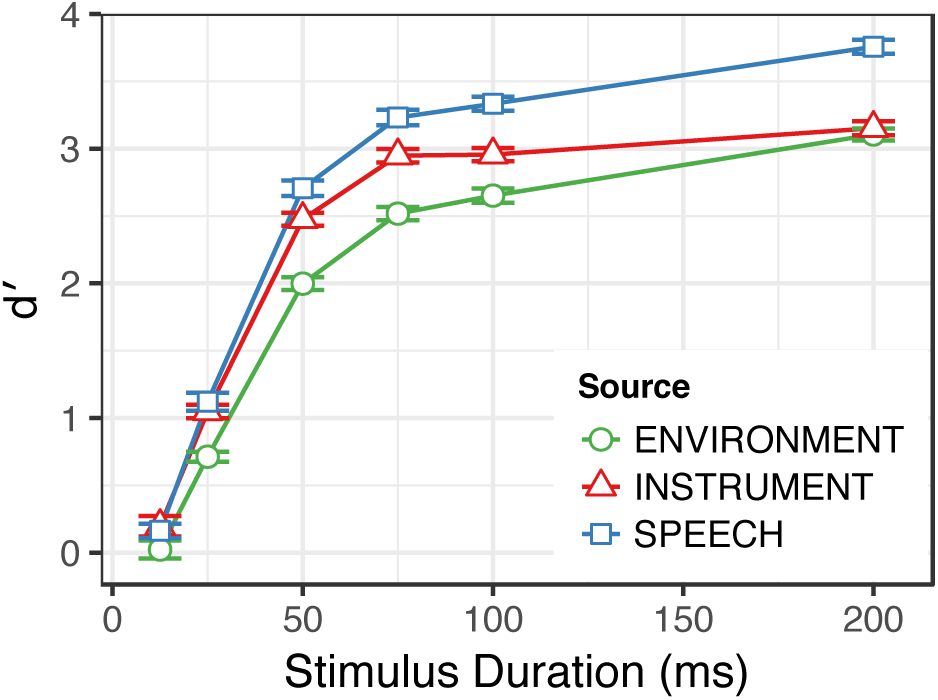
*d′* scores of participant responses in each block of Experiment 1 displayed by target sound-class as a function of gate duration.

Beyond the class-level results, Figure 2 depicts a number of interesting confusion patterns for specific items. Vowels were correctly classified very accurately very early on (by 25-ms per Figure 2), while consonants were frequently mis-classified as human-environmental sounds. Somewhat surprisingly, a few instrument sounds were mis-classified as human-environmental sounds. These instrument tokens tended to exhibit a rough quality either in the hammered excitation of the harpsichord notes, or via some harshness in the bowed attack of one particular cello arco token. Some human-environmental sounds were also mis-classified, specifically some air inflation/deflation sounds (both exhibiting a bright, noisy sibilant quality similar to the /ʃ/ in the speech tokens) as well as some higher-frequency and more pitched impact sounds (exhibiting a bright, rough percussive quality similar to the harpsichord).

Together, these figures indicate that, (as expected) participant performance improved as duration increased through successive blocks. Furthermore, while there appeared to be a bias towards making human environmental responses at early gates (high accuracy, low *d’*; see also Figure 3). Participants were more discerning in their speech (low accuracy early, but high *d’*) and instrument responses (high accuracy, high *d’*) with an advantage for speech stimuli emerging across all measures at the longest gates.

Classification performance significantly exceeded chance (*d’* = 0) for all categories by the 25-millisecond gate (one-sample *t*-tests, *df* = 85, speech *t* = 13.2; instrument *t* = 14.5; human-environmental *t* = 15.4; all two-tailed *p* < 0.001, significant against a Bonferroni correction across the six gates: *p* < 0.0083). Performance at the 12.5 millisecond gate exceeded chance for speech (*t* = 4.0, *p* < 0.001) and instrument (*t* = 2.9, *p* = 0.005) responses but not for human-environmental responses (*t* = 0.5, *p* = 0.595).

Mixed-effect models of class responses fit to trial-level accuracy (Table 2) and block-level *d’* scores (Table 3), both revealed significant main effects of duration, and significant (negative) interactions between duration and the instrument and environment response categories relative to the speech class intercept. Musical training interacted to improve trial-level accuracy performance at later duration gates (significant positive interaction between musical training and duration), particularly for human-environmental stimuli (significant positive interaction between musical training and human-environmental sound accuracy relative to speech sounds), and these effects were moderated by negative interactions for musical instrument stimuli (negative two- and three-way interactions between instrument stimuli, musical training and duration). However, these musical training results for accuracy should be interpreted with caution since the *d’* analyses suggest a substantially reduced influence of musical training when false alarms are taken into account (Table 3), suggesting that musicians perhaps used a different strategy for responding, but that such a strategy was not necessarily more discriminating. Speech *d’* scores were also better than both instrument and human-environmental *d’* scores across duration gates (significant main effects).

**Table 2.** Results of the Generalized Binomial Mixed-Effects Model of Trial-Level Accuracy.

| Parameters | b | SE | z | p |
| --- | --- | --- | --- | --- |
| (Intercept) | 1.34 | 0.12 | 10.95 | < 0.001 |
| Human-Environment | 0.07 | 0.15 | 0.48 | 0.632 |
| Instrument | 0.18 | 0.15 | 1.19 | 0.233 |
| Duration | 1.54 | 0.07 | 22.87 | < 0.001 |
| Musical Training | 0.08 | 0.06 | 1.33 | 0.185 |
| Duration * Human-Environment | -0.45 | 0.08 | -5.66 | < 0.001 |
| Duration * Instrument | -0.30 | 0.08 | -3.76 | < 0.001 |
| Musical Training * Human-Environment | 0.12 | 0.03 | 4.06 | < 0.001 |
| Musical Training * Instrument | -0.08 | 0.03 | -2.58 | 0.010 |
| Duration * Musical Training | 0.10 | 0.04 | 2.48 | 0.013 |
| Duration * Musical Training * Human-Environment | -0.05 | 0.03 | -1.62 | 0.106 |
| Duration * Musical Training * Instrument | -0.14 | 0.03 | -4.74 | < 0.001 |

Formula: (Accuracy) ~ (Target-Class) \* (Duration) \* (Musical Training) + (Duration||Subject) + (Duration||Item)
Target-class: dummy coded with Speech as the reference level.
Duration: log-transformed and z-normalized
Musical Training: Gold-MSI subscale, z-normalized

**Table 3.** Results of the Linear Mixed-Effects Model of Participant’s Block-Level d′ Scores.

| Parameters | b | SE | t | p |
| --- | --- | --- | --- | --- |
| (Intercept) | 2.39 | 0.06 | 43.32 | < 0.001 |
| Human-Environment | -0.55 | 0.03 | -16.04 | < 0.001 |
| Instrument | -0.26 | 0.03 | -7.45 | < 0.001 |
| Duration | 1.26 | 0.03 | 38.56 | < 0.001 |
| Musical Training | 0.04 | 0.06 | 0.79 | 0.433 |
| Duration * Human-Environment | -0.18 | 0.03 | -5.17 | < 0.001 |
| Duration * Instrument | -0.20 | 0.03 | -5.93 | < 0.001 |
| Musical Training * Human-Environment | 0.06 | 0.03 | 1.74 | 0.081 |
| Musical Training * Instrument | 0.04 | 0.03 | 1.16 | 0.245 |
| Duration * Musical Training | -0.03 | 0.03 | -1.05 | 0.297 |
| Duration * Musical Training * Human-Environment | 0.07 | 0.03 | 2.05 | 0.041 |
| Duration * Musical Training * Instrument | 0.04 | 0.03 | 1.18 | 0.238 |

Formula: $(d') \sim (\text{Target-Classs}) * (\text{Duration}) * (\text{Musical Training}) + (\text{Duration}||\text{Subject})$
Target-class: dummy coded with Speech as the reference level.
Duration: log-transformed and z-normalized
Musical Training: Gold-MSI subscale, z-normalized

Participant responses for each class (at the trial-level) were also modeled as a function of the acoustic features of the stimuli (on each trial). To determine the most parsimonious set of acoustic features associated with participant responses for each class the data were analyzed via an all-subsets regression analysis (run for each sound-class separately; see Section II-A.5 for details). The results of these analyses are summarized in Table 4. The full models of these acoustic features and their interactions with duration can be found in Supplemental Table 1.

**Table 4.** Model Comparison Analysis of Trial-level Class Responses: Averaged Model Results Within 3 BIC of Top.

| Model Predictor | <u>Instrument</u> |  |  |  | <u>Human-Environment</u> |  |  |  | <u>Speech</u> |  |  |  |
| --- | --- | --- | --- | --- | --- | --- | --- | --- | --- | --- | --- | --- |
|  | Relative Importance | b | SE | p | Relative Importance | b | SE | p | Relative Importance | b | SE | p |
| Aperiodicity | 1 | -0.25 | 0.03 | < 0.001 | 1 | 0.21 | 0.03 | < 0.001 |  |  |  |  |
| Log-Attack-Time | 1 | 0.60 | 0.11 | < 0.001 |  |  |  |  | 1 | -0.57 | 0.17 | < 0.001 |
| Spectral Centroid – IQR | 0.79 | -0.09 | 0.02 | < 0.001 |  |  |  |  | 1 | 0.24 | 0.03 | < 0.001 |
| Spectral Centroid – Median | 1 | -0.62 | 0.08 | < 0.001 | 1 | 0.37 | 0.06 | < 0.001 |  |  |  |  |
| Spectral Flatness – IQR |  |  |  |  | 1 | -0.32 | 0.03 | < 0.001 | 1 | 0.21 | 0.04 | < 0.001 |
| Spectral Flatness – Median |  |  |  |  | 1 | 0.34 | 0.04 | < 0.001 | 1 | -0.48 | 0.06 | < 0.001 |
| Spectral Variability – Median |  |  |  |  |  |  |  |  | 0.79 | -0.15 | 0.03 | < 0.001 |
| Temporal Centroid |  |  |  |  | 1 | -3.72 | 0.27 | < 0.001 |  |  |  |  |
| Duration | 1 | -1.12 | 0.13 | < 0.001 | 1 | 3.14 | 0.26 | < 0.001 | 0.21 | 0.41 | 0.23 | < 0.001 |
| Duration * Spectral Centroid – IQR |  |  |  |  |  |  |  |  | 0.21 | 0.12 | 0.02 | < 0.001 |
| Duration * Spectral Centroid – Median | 1 | -0.29 | 0.04 | < 0.001 |  |  |  |  |  |  |  |  |
| Duration * Spectral Flatness – IQR |  |  |  |  | 1 | -0.16 | 0.02 | < 0.001 |  |  |  |  |
| Duration * Temporal Centroid |  |  |  |  | 1 | 1.11 | 0.09 | < 0.001 |  |  |  |  |

Formula: Varied depending on best fitting models in Stage 1 of the all subsets regression
One model was fit to the data for each binary class identification.
All acoustic features were z-normalized in the model.
Duration: log-transformed and z-normalized.

These findings generally implicate the same features that influenced responses for each class in previous work (Ogg et al., 2017) with a few additional features implicated in these results. Musical instrument responses were again characterized by low aperiodicity and low, less variable spectral centroids. However, instrument responses were also associated with longer onset times (positive main effect of log attack time) and an additional influence of lower spectral centroids at later durations (negative interaction between spectral centroid and duration). Human environmental responses were again characterized by high aperiodicity and high spectral centroids. These sounds were also characterized by higher, less variable spectral flatness values, and earlier temporal centroids. Speech responses were again characterized by more variable spectral centroids (particularly at later gates). Speech responses were also associated with lower but more variable spectral flatness, shorter onsets and lower spectral variability overall. Thus, a largely similar picture of results was obtained using this new set of stimuli and three-choice task as in prior work using a gating paradigm (Ogg et al., 2017; also similar to a related dissimilarity rating task, Ogg & Slevc, 2019a). This correspondence between the analyses of the natural sounds and previous work indicates that the synthesized stimulus manipulations are also generally applicable for these data, allowing for a stronger test of the influence of these acoustic qualities on classification behavior.

#### 2. Analyses of synthesized sound tokens

Sound-class response rates for the synthesized stimuli are summarized across duration gates in Figure 6. In general, it is clear that the aperiodicity manipulation had a large influence on human-environmental (associated with high aperiodicity, i.e., the noise carrier) and instrument responses (associated with low aperiodicity, i.e., the sawtooth wave carrier). It is also clear that the rate of speech responses for the synthesized stimuli dropped off sharply throughout the earliest gates. The influence of the other manipulations for spectral centroid, spectral variability (both which interacted with the aperiodicity manipulation) and fundamental frequency appear to be subtle compared with the aperiodicity manipulation.

**Figure 6.**
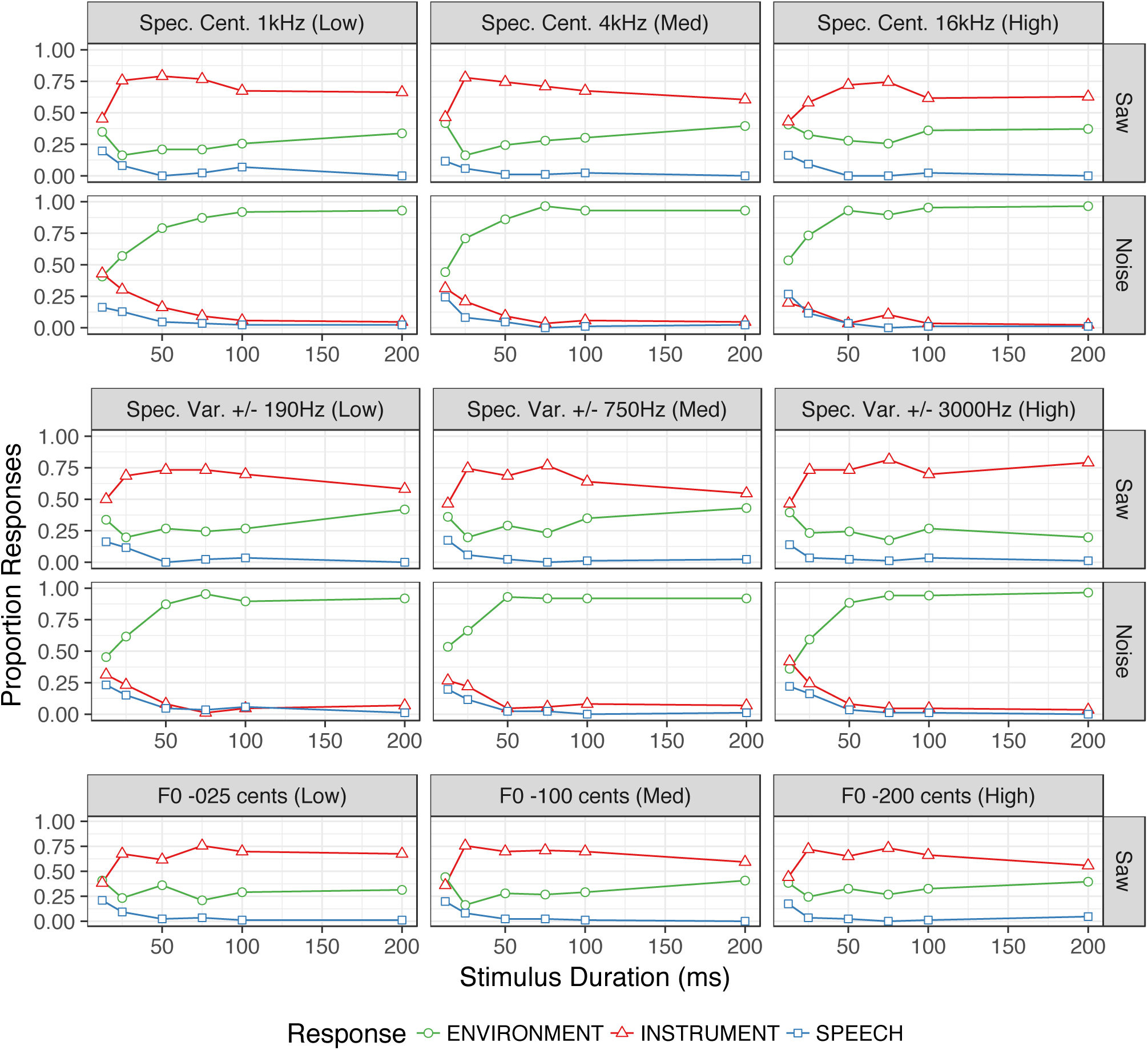
Summary of class responses for synthesized stimuli in Experiment 1 across durations by acoustic manipulation condition. Predictions are plotted as a function of the acoustic manipulation conditions (same layout as Figure 1) and averaged over participants.

To examine the influence of these manipulations on responses for each class, a series of binomial mixed-effect regressions was fit to each binary class response and manipulation (i.e., the three-choice responses for each trial “one-hot” encoded and analyzed for the subset of trials corresponding to each manipulation). Binary class responses on each of these trials were then analyzed as a function of variables coding the experimental conditions (duration and aperiodicity crossed with the spectral centroid, variability or fundamental frequency manipulation). These models are summarized in Table 5.

**Table 5.** Experiment 1, Models of Participant Class Responses to Synthesized Stimulus Manipulations.

| Model Predictor | Fundamental Frequency Manipulation |  |  |  |  |  |  |  |  |  |  |  |
| --- | --- | --- | --- | --- | --- | --- | --- | --- | --- | --- | --- | --- |
|  | Instrument |  |  |  | Human-Environment |  |  |  | Speech |  |  |  |
|  | b | SE | z | p | b | SE | z | p | b | SE | z | p |
| (Intercept) | 0.83 | 0.18 | 4.52 | < 0.001 *** | -1.22 | 0.19 | -6.28 | < 0.001 *** | -3.45 | 0.31 | -11.21 | < 0.001 *** |
| Fundamental Frequency -1 cents (Med) | -0.01 | 0.15 | -0.07 | 0.948 | 0.05 | 0.16 | 0.33 | 0.744 | -0.34 | 0.44 | -0.78 | 0.438 |
| Fundamental Frequency -2 cents (High) | -0.07 | 0.15 | -0.49 | 0.624 | 0.16 | 0.16 | 0.99 | 0.321 | -0.09 | 0.39 | -0.23 | 0.819 |
| Duration | 0.58 | 0.14 | 4.14 | < 0.001 *** | -0.20 | 0.14 | -1.41 | 0.158 | -1.14 | 0.24 | -4.71 | < 0.001 *** |
| Fundamental Frequency -1 cents (Med) * Duration | -0.18 | 0.15 | -1.18 | 0.238 | 0.19 | 0.16 | 1.21 | 0.228 | -0.19 | 0.35 | -0.54 | 0.591 |
| Fundamental Frequency -2 cents (High) * Duration | -0.36 | 0.15 | -2.32 | 0.020 * | 0.23 | 0.16 | 1.44 | 0.149 | 0.31 | 0.33 | 0.94 | 0.348 |
|  | Spectral Centroid Manipulation |  |  |  |  |  |  |  |  |  |  |  |
|  | Instrument |  |  |  | Human-Environment |  |  |  | Speech |  |  |  |
|  | b | SE | z | p | b | SE | z | p | b | SE | z | p |
| (Intercept) | -0.60 | 0.15 | -4.11 | < 0.001 *** | 0.17 | 0.14 | 1.23 | 0.218 | -3.31 | 0.21 | -15.85 | < 0.001 *** |
| Noise | -3.17 | 0.20 | -16.06 | < 0.001 *** | 3.05 | 0.19 | 16.27 | < 0.001 *** | 0.28 | 0.34 | 0.82 | 0.410 |
| Spec. Cent. 4kHz (Med) | -0.32 | 0.14 | -2.32 | 0.020 * | 0.41 | 0.13 | 3.14 | 0.002 ** | -0.55 | 0.29 | -1.90 | 0.057 . |
| Spec. Cent. 16kHz (High) | -0.59 | 0.14 | -4.10 | 0.000 *** | 0.63 | 0.13 | 4.67 | < 0.001 *** | -0.68 | 0.31 | -2.17 | 0.030 * |
| Duration | -0.45 | 0.11 | -4.13 | 0.000 *** | 0.69 | 0.11 | 6.14 | < 0.001 *** | -0.95 | 0.15 | -6.16 | < 0.001 *** |
| Noise * Spec. Cent. 4kHz (Med) | -0.37 | 0.28 | -1.32 | 0.188 | 0.23 | 0.26 | 0.87 | 0.386 | 0.27 | 0.58 | 0.47 | 0.639 |
| Noise * Spec. Cent. 16kHz (High) | -0.47 | 0.29 | -1.63 | 0.104 | 0.29 | 0.27 | 1.07 | 0.283 | 0.24 | 0.62 | 0.39 | 0.696 |
| Noise * Duration | -1.56 | 0.18 | -8.46 | < 0.001 *** | 1.44 | 0.18 | 7.93 | < 0.001 *** | 0.20 | 0.30 | 0.66 | 0.508 |
| Spec. Cent. 4kHz (Med) * Duration | -0.02 | 0.13 | -0.16 | 0.874 | 0.06 | 0.13 | 0.45 | 0.656 | -0.28 | 0.24 | -1.18 | 0.239 |
| Spec. Cent. 16kHz (High) * Duration | 0.18 | 0.13 | 1.37 | 0.170 | -0.03 | 0.13 | -0.20 | 0.843 | -0.60 | 0.25 | -2.40 | 0.016 * |
| Noise * Spec. Cent. 4kHz (Med) * Duration | 0.26 | 0.26 | 1.00 | 0.318 | 0.02 | 0.26 | 0.08 | 0.934 | -0.39 | 0.48 | -0.82 | 0.414 |
| Noise * Spec. Cent. 16kHz (High) * Duration | 0.29 | 0.27 | 1.07 | 0.284 | 0.08 | 0.26 | 0.30 | 0.765 | -0.26 | 0.50 | -0.52 | 0.604 |
|  | Spectral Variability Manipulation |  |  |  |  |  |  |  |  |  |  |  |
|  | Instrument |  |  |  | Human-Environment |  |  |  | Speech |  |  |  |
|  | b | SE | z | p | b | SE | z | p | b | SE | z | p |
| (Intercept) | -0.92 | 0.15 | -6.28 | < 0.001 *** | 0.42 | 0.14 | 3.00 | 0.003 ** | -3.25 | 0.21 | -15.69 | < 0.001 *** |
| Noise | -3.41 | 0.21 | -15.97 | < 0.001 *** | 3.04 | 0.19 | 15.98 | < 0.001 *** | 0.71 | 0.36 | 1.96 | 0.050 . |
| Spec. Var. +/- 750Hz (Med) | 0.02 | 0.14 | 0.11 | 0.912 | 0.15 | 0.13 | 1.19 | 0.236 | -0.47 | 0.29 | -1.63 | 0.103 |
| Spec. Var. +/- 3000Hz (High) | 0.18 | 0.15 | 1.19 | 0.234 | -0.02 | 0.14 | -0.14 | 0.891 | -0.37 | 0.28 | -1.35 | 0.179 |
| Duration | -0.45 | 0.11 | -4.23 | < 0.001 *** | 0.73 | 0.10 | 6.97 | < 0.001 *** | -1.07 | 0.16 | -6.80 | 0.000 *** |
| Noise * Spec. Var. +/- 750Hz (Med) | 0.19 | 0.28 | 0.68 | 0.500 | 0.05 | 0.26 | 0.20 | 0.840 | -0.54 | 0.58 | -0.95 | 0.345 |
| Noise * Spec. Var. +/- 3000Hz (High) | -0.33 | 0.30 | -1.11 | 0.267 | 0.51 | 0.27 | 1.86 | 0.063 . | -0.52 | 0.55 | -0.95 | 0.344 |
| Noise * Duration | -1.27 | 0.19 | -6.59 | < 0.001 *** | 1.22 | 0.18 | 6.75 | < 0.001 *** | 0.18 | 0.30 | 0.58 | 0.561 |
| Spec. Var. +/- 750Hz (Med) * Duration | 0.09 | 0.13 | 0.68 | 0.499 | -0.05 | 0.12 | -0.43 | 0.668 | -0.18 | 0.24 | -0.78 | 0.434 |
| Spec. Var. +/- 3000Hz (High) * Duration | 0.04 | 0.14 | 0.27 | 0.790 | -0.02 | 0.13 | -0.13 | 0.900 | -0.10 | 0.23 | -0.45 | 0.651 |
| Noise * Spec. Var. +/- 750Hz (Med) * Duration | 0.31 | 0.26 | 1.19 | 0.234 | -0.17 | 0.25 | -0.67 | 0.503 | -0.32 | 0.47 | -0.68 | 0.496 |
| Noise * Spec. Var. +/- 3000Hz (High) * Duration | -0.68 | 0.27 | -2.51 | 0.012 * | 0.99 | 0.26 | 3.73 | < 0.001 *** | -0.72 | 0.46 | -1.56 | 0.120 |

Formula: ('Class Present' Response) ~ (Noise) \* (Manipulation) \* (Duration) + (Duration|Subject)
One model was fit to the data for each binary class response variable (column sections) and manipulation condition (row sections).
Noise: 2 Contrast Coded Levels ("Noise", "Saw") "Saw" as reference. Note this predictor was not present in the model of the Fundamental Frequency Manipulation.
Manipulation (broken down by subheadings): 3 Dummy Coded Levels ("Low", "Medium", "High") with "Low" as the reference level.
Duration: log-transformed and z-normalized

In support of the observations above, large contrasting effects were observed for the aperiodicity manipulation across all conditions for instrument (significantly more responses to the low aperiodicity, saw tooth wave carrier) and human environmental responses (significantly more responses to the high aperiodicity, noise carrier), although speech responses in all conditions decreased as duration increased. The other manipulations had a smaller influence but some main effects are notable. First, higher fundamental frequency variability reduced instrument responses at longer durations. Second, the spectral centroid manipulation had a similar (albeit weaker) contrasting effect as the aperiodicity manipulation on music (reduced response rates for the higher spectral centroids) and human-environmental (increased response rates for the higher spectral centroids) responses. Fewer speech responses were also observed in the highest spectral centroid condition, especially at later durations. Finally, the effects of spectral variability were quite limited finding that the largest spectral variability manipulation (i.e., the largest low-pass filter sweep) interacted with the noise condition at longer durations (three-way interaction), resulting in more human-environmental responses (for the noise stimulus with high variability; relative to the sawtooth wave as the intercept) and fewer instrument responses.

Thus, while speech stimuli were difficult to characterize in this synthesis approach, the pattern of effects observed for the spectral centroid and aperiodicity manipulations with respect to the human environmental and instrument sounds aligned with the acoustic modeling analyses of the natural sounds. The effects observed for the fundamental frequency and spectral variability manipulations were more subtle but at least partially aligned with hypotheses for instrument sounds and environmental sounds suggested by past work.

### C. Discussion

This study examined three-choice sound classification responses for duration-gated stimuli. The experiment was designed to test prior results implicating acoustic qualities related to noisiness, spectral envelope, spectrotemporal variability, and fundamental frequency variability (Gygi et al., 2007; Huang & Elhilali, 2017; Ogg et al., 2017; Ogg & Slevc 2019a; Ogg et al., 2020) in how listeners distinguish speech, musical instrument, and human-environmental sounds. In addition to a novel set of natural sounds, a set of synthesized stimuli were presented that were designed to manipulate the acoustic dimensions of interest. The results of this study broadly aligned with the most closely related previous work (Ogg et al., 2017), despite key differences related to the stimulus set (different exemplars/recordings of speech, musical instrument, or human-environmental sounds) and task (a three-choice classification task rather than a go/no-go task). This experiment also involved a much larger participant sample than in prior work. Analyses of the classification responses for the synthesized stimuli converged with many of the main hypotheses. The results for the synthesized stimuli also broadly aligned with previous work: acoustic models of class-responses for natural stimuli (see also Ogg et al., 2017), dissimilarity ratings (Ogg & Slevc 2019a) and electrophysiological (Ogg et al., 2020) results. Together this provides additional support for the notion that shortly after sound onset listeners use noise and spectral envelope (centroid) cues and to a lesser degree spectral variability, and fundamental frequency cues in making judgments about the kind of sounds they hear. Moreover, the clearest effects are for aperiodicity and spectral centroid with respect to instrument and human-environmental sound classification.

There were, however, a number of ways in which these results diverged from prior work which warrant discussion. First, and most notably, participants in this study were more accurate at discriminating speech sounds from the other sound classes across time points, relative to Ogg and colleagues (2017; but in line with Agus et al., 2012 and Suied et al., 2014), evidenced by significantly higher *d’* for speech responses than other sound class responses across durations. This is attributable to differences in the studies’ stimulus sets. Ogg and colleagues (2017; Experiment 1) found that speech consonants were frequently confused with human-environmental sounds, which was also observed here (Figure 2). However, the vowels that were included here were very accurately classified as speech even at early gates (Figure 2), suggesting a perceptual benefit for this variety of speech sounds similar to previous findings (Agus et al., 2012; Isnard et al., 2019; Suied et al., 2014). Performance on these vowels thus improved overall accuracy for speech responses.

At the same time, however, the speech stimuli did not appear to be well characterized by the synthesized stimulus manipulations. Indeed, the most consistent finding regarding speech responses for the synthesized sounds was for participants to make very fewer speech responses as the synthesized stimuli increased in duration. Speech responses were not influenced by increased spectral or fundamental frequency variability among the synthesized stimuli as hypothesized. Given participants’ especially accurate performance for vowel stimuli, perhaps a different, more speech-specific synthesis approach (that incorporated formants or other voicing cues, for example) might have elicited more responses. Unfortunately, such a manipulation was not immediately apparent from the acoustic features that were implicated in prior work. Thus, this issue would be well suited for examination in future studies perhaps via morphing or other methods.

Acoustic features associated with classification responses for speech, musical instrument and human-environmental sounds exhibited a large degree of overlap with Ogg and colleagues (2017). Instrument and environmental responses were characterized by contrasting values for spectral centroid and aperiodicity, while speech stimuli were strongly influenced by specific spectral changes over time (variable centroid and spectral flatness). These findings for instrument and environmental responses also aligned classification of the synthesized sounds. However, temporal cues played a larger role in these data than previous gating studies (Ogg et al., 2017). Temporal envelope cues were associated with responses in all categories: onset velocity, characterized by log-attack-time was related to instrument and speech responses and the temporal centroid was a cue for environmental responses. Because previous work did not suggest a strong influence of temporal cues, the temporal envelopes of the synthesized stimuli were held constant and not manipulated among the synthesized stimuli. However, this would be an excellent avenue for future research. This is also true for stimulus offsets or decays, which were not examined in favor of an emphasis on early object recognition processes (i.e., around onset). There also appeared to be a stronger influence of spectral flatness for speech and human-environmental sounds than previously observed. Spectral flatness is a measure that is similar to noisiness but based on the frequency spectrum and is thus related to the aperiodicity manipulation (which generally aligns with the acoustic modeling results for the human-environmental sounds).

Finally, rather nuanced results were obtained for the spectral variability manipulation among the synthesized stimuli. Previous findings suggested a strong influence of this dimension (Gygi et al., 2007; Huang & Elhilali, 2017; Ogg & Slevc, 2019a; Ogg et al., 2017), but it is also a potentially complex and multifaceted acoustic attribute, which was given a rather general treatment here (c.f., Thoret et al., 2016; Venezia et al., 2016). The specific variability manipulation used here essentially introduces variability into the spectral centroid feature. Participants were more likely to make more human-environmental responses (and fewer instrument responses) to the high spectral variability manipulation applied to the noise carrier when these stimuli were given time to develop at later duration gates. Contrary to the importance of spectral centroid IQR for speech stimuli in the models of classification responses for the natural sounds, the spectral variability manipulation did not influence rates of speech responses. In addition to the possibility these stimuli were more or less discounted by participants as patently “not speech” (see discussion above), closer examination of the synthesized high spectral variability stimuli suggests that, when applied to the sawtooth wave sounds the filter sweep sounds somewhat harsh and closely resembles the opening transient (or “blat”) of a brass instrument onset. Thus, this manipulation did not characterize overall spectral variability as well as some other manipulations might have. Perhaps a less predictable trajectory for the movement of the filter cutoff (e.g., one controlled by a random noise signal) rather than the up/down ramp used here would be more similar to a human environmental or speech sound. In line with this suggestion, it seems that participants were more likely to make more human-environmental responses (and fewer instrument responses) to the high spectral variability manipulation applied to the noise carrier when these stimuli were given time to develop at later duration gates.

Thus, these results generally align with the main hypotheses and with previous studies, albeit with some nuance in in terms of class accuracy (greater accuracy for speech stimuli overall) and some of the specific acoustic features that influenced task performance. These differences might be due to variations in stimulus sets and speech stimuli in particular (i.e., the inclusion of both consonants and vowels), as well as the task (three-choice, vs go/no-go or item-level identification and ratings), or the sample size. Importantly, however, despite these ostensible differences, the acoustic modeling results generally implicated a similar set of acoustic cues used differentiate sound classes and analyses of the synthesized sounds converged on many of the same key acoustic attributes. Indeed, the similar acoustic modeling results for the natural stimuli (especially the instrument and environmental sounds) in this study provide a solid grounding for interpretations of the synthesized stimuli, given that the synthesized sounds were designed to manipulate the same acoustic features implicated in previously. Together, these results indicate that cues relating to aperiodicity, spectral centroid, (and perhaps to a lesser degree) spectral variability, and fundamental frequency appear to support listeners’ ability to differentiate sounds of different super-ordinate classes. However, to more directly examine the processes by which listeners and the auditory system transform this acoustic information into representations of sounds and categories, it would be useful to examine the neural processes that rapidly construct these representations after sound onset. Such an investigation requires the collection of temporally resolved neural responses, which was undertaken in Experiment 2.

## III. EXPERIMENT 2: EEG DECODING STUDY

Understanding how the brain accumulates acoustic information in service of sound recognition requires delineating the neural computations that take place as perception unfolds alongside the developing acoustic events. While gated classification tasks like Experiment 1 can provide part of the answer, behavioral measures are a summation of the bottom-up and top-down processing that occurs between auditory perception and the execution of a motor action to register a response. To provide a clearer picture of what processes facilitate rapid auditory object and event recognition we must understand how neural representations evolve immediately following sound onset. Previous M/EEG studies of auditory object and event recognition (e.g., Zuk et al., 2020) provide some insight and broadly complement behavioral results: neural responses appear to differ by superordinate sound-classes as early as 70 (in EEG: Charest et al., 2009; Murray et al., 2006) or 90 (in MEG: Lowe et al., 2020; Ogg et al., 2020) milliseconds after onset. Some studies suggest a stronger association with acoustic features prior to 200 milliseconds (Ogg et al., 2020), with more semantic or categorical representations dominating afterwards (Lowe et al., 2020). However, a more thorough and guided search of this acoustic feature space is needed since not all previous studies clarify what acoustic features might influence these observed acoustic or class-wise responses beyond high level acoustic controls (Charest et al., 2009; Lowe et al., 2020; Murray et al., 2006) or manipulations (Zuk et al., 2020). Ogg and colleagues (2020) provide some insight and implicated a feature set that largely aligns with the behavioral data in Experiment 1 (i.e., aperiodicity, spectral envelope and spectrotemporal change over time). However, these findings were correlational and did not involve and manipulations of the features or dimensions of interest to better understand their relative importance.

Experiment 2 aims to address some of these critiques using the high temporal resolution of EEG. The stimuli comprised the same set of synthesized sounds from Experiment 1 along with a subset of that study’s natural sounds (see Figure 7). Again, in addition to testing previous MEG results (Ogg et al., 2020) in a new imaging modality (EEG) and with new stimuli, the expectation was that the synthesized manipulations might better isolate the influence of specific acoustic dimensions. Synthesized manipulations of these acoustic parameters were hypothesized to influence participants’ neural responses in a way that might resemble responses to different classes of the natural sounds. The degree of correspondence between the natural sound classes and acoustic feature manipulations was assessed by training a pattern classifier on the superordinate sound-class labels associated with neural responses to the natural sound set and “testing” the classifier on responses to the synthesized stimuli (i.e., eliciting classification hypothesis: speech, instrument or human-environment). Since the synthesized stimuli are simple distillations (and manipulations) of acoustic features hypothesized to play a role in neural representations of each sound class, the classifier’s “predictions” for the synthesized sounds in this paradigm yield an indication of how closely the neural responses throughout an acoustic dimension resemble responses to natural sounds within each class.

**Figure 7.**
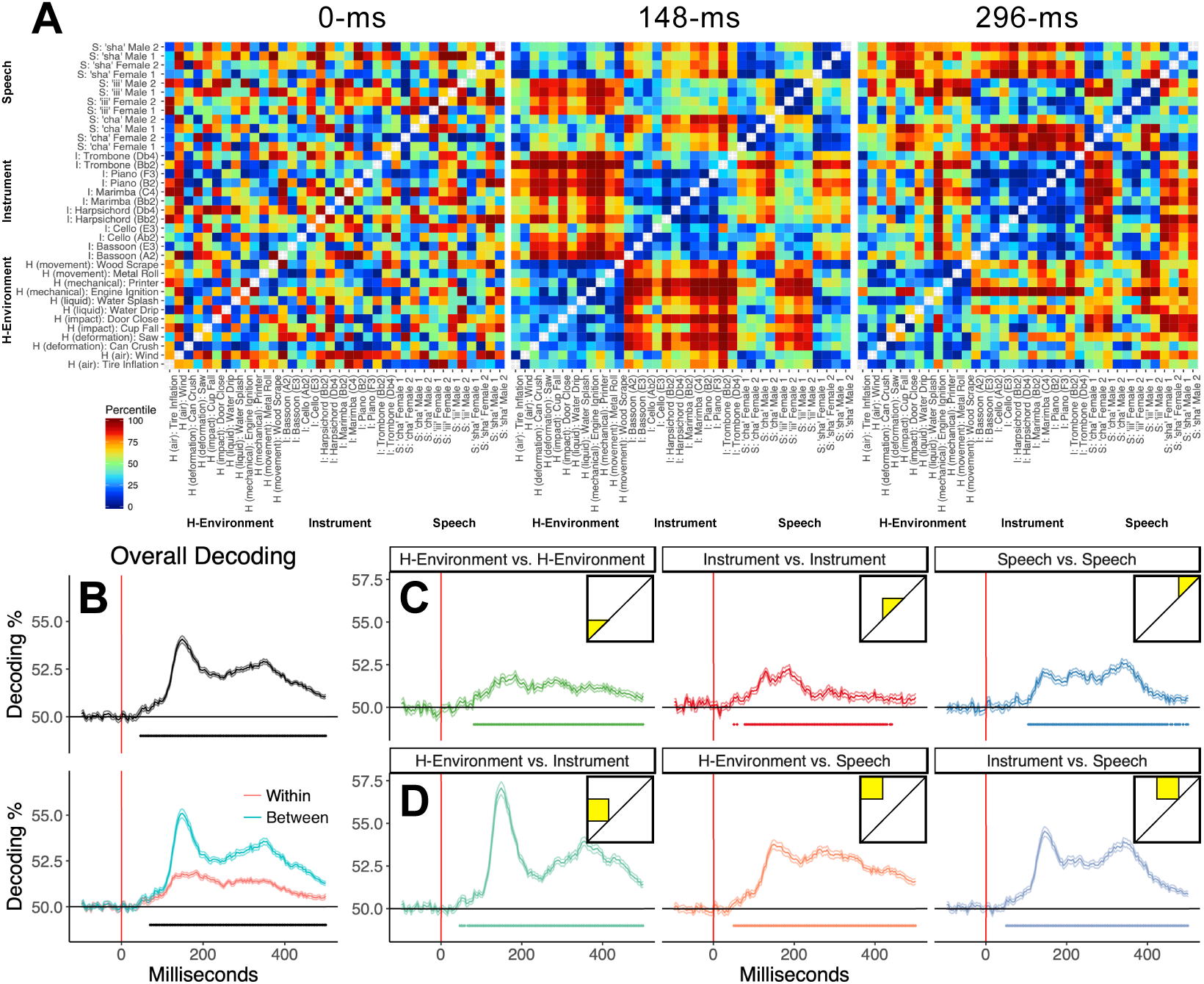
Decoding accuracy for natural auditory objects and events in Experiment 2. (A) Group-averaged neural dissimilarity matrices (RDMs) at 0, 148, and 296 ms. (B) Grand average decoding accuracy (top; * *p* < 0.05 corrected v.s. chance) and average between v.s. within-class decoding accuracy (bottom; * *p* < 0.05 corrected between v.s. within). (C) Within-class decoding accuracy for each sound class (* *p* < 0.05 corrected v.s. chance). (D) Between-class decoding accuracy for each pair of sound categories (* *p* < 0.05 corrected v.s. chance). Square insets depict the region of the RDM that was used for each comparison. The top of B encompasses all the insets in C and D, and the bottom of B represents the insets of C v.s. D. Lines indicate +/– one standard error. Note that RDMs are symmetrical across the diagonal so only the upper half of the matrix was analyzed, although both sides of the diagonal are depicted in this figure.

A notable departure of Experiment 2 from the prior work it was based on (and is partially re-examining: Ogg et al., 2020) is in the use of EEG rather than MEG. Relative to MEG, EEG is comparatively cheap, portable, widely available, and sensitive to both tangential and radial neural sources (Ahlfors, Han, Belliveau, & Hämäläinen, 2010; Baillet, 2017). However, compared with MEG, EEG suffers a loss in spatial resolution that will likely influence overall decoding performance in two ways. First, this EEG set up employed a reduced number of sensors around the scalp relative to most modern MEG systems, which limits the features available for training and testing the classifier (64 EEG channels here vs. 157 MEG channels previously). Second, EEG is subject to distortions of the neural signal that are incurred by the skull and other tissues (Baillet, 2017). On the other hand, the audio playback apparatus in the MEG study reduced high frequency acoustic information (< approximately 5.5 kHz) during stimulus playback, whereas the EEG set up was not limited in this way. Given these various constraints, Experiment 2 represents an interesting test and complement to prior work.

### **A.** Methods and materials

#### 1. Participants

Forty-two new participants were recruited from the University of Maryland’s Psychology Department (30 female, 4 left handed, age: M = 21, SD = 4.1, min = 18, max = 40). Participants were compensated with money or course credit. All participants reported normal hearing. Recruitment was carried out irrespective of musical ability, so most participants (79%) had obtained some degree of musical training (M = 5.4, SD = 3), similar to Experiment 1. EEG data from ten other participants (not counted among the forty-two above) were removed prior to further analysis: two because they did not complete all four runs, one because they fell asleep during the study, two for excessive noise in their EEG, and five participants’ data were excluded because of an errant trigger issue. These last participants’ data were removed out of an abundance of caution, but including them (following a fix for the trigger issue) yielded the same group-level results.

#### 2. Stimuli and apparatus

Experiment 2 presented participants with 36, 300 millisecond sound tokens that were a representative subset of the stimuli from Experiment 1 (and different from Ogg et al., 2020). These stimuli were evenly divided among 12 speech (/tʃa/, /ʃa/, and /i/ utterances from 2 male and 2 female speakers), 12 musical instrument (marimba, trombone, bassoon, cello arco, piano, harpsichord each playing 2 notes corresponding to the fundamental frequency of one male and one female utterance) and 12 human-environmental sound tokens (2 sounds representing each of the following excitation methods/media: air, deformation, impact, mechanical, movement, and liquid). The same 15 synthesized stimuli from Experiment 1 (see Section II-A.2) were also added to this stimulus set. The only difference in pre-processing for Experiment 2 was that all stimuli now had a 300-millisecond duration from onset. A dog vocalization was also included as a catch trial to confirm that participants attended to the sounds during the EEG session. All sound tokens were normalized for their root-mean square level and were presented via a Mackie CR5BT speaker positioned at approximately eye level 42 inches in front of the participant during the EEG session. This achieved an average playback level of approximately 65dB-A.

#### 3. Procedure

The procedure for Experiment 2 was similar to Ogg and colleagues 2020 with minor adjustments owing to the EEG set up and the higher fidelity of the stimulus presentation system. Before participating in the EEG session, participants attended a short (half-hour) screening and familiarization visit within 14 days (M = 5.7, SD = 2.9) of the EEG session. At the screening visit, participants first listened freely to the labeled stimulus sound files as many times and in whatever order they liked prior to starting the screening protocol (in PsychoPy version 1.83.4; Peirce, 2007). During the screening task, participants were presented with three blocks of trials. Stimuli were presented three times on each trial with a 700-millisecond inter-stimulus-interval. The first block of trials simply presented the stimuli in a random order with their labels (from the response key, same as the sound file names) displayed on the screen. No responses were required during the first block. The second block of trials presented the stimuli in a random order and at the end of each trial the participant was asked to indicate which sound they heard via a button press from a response key. Note, for speech tokens, participants needed to identify both the speaker (each assigned novel, gender consistent names) and the speech utterance. After each response in the second block, participants received feedback to indicate if they were correct or to indicate the correct sound and response if they were incorrect. The third block was the same as the second block, but no feedback was provided.

Participants were required to achieve an accuracy criterion of over 80% on the third block of trials in order to proceed to the EEG session. If a participant did not reach this criterion, they were allowed to freely re-listen to the stimuli and retry the third screening block (performance at criterion M = 88.7%, SD = 5%). Note, this is a slightly lower accuracy at criterion than in previous work (Ogg et al., 2019; Ogg et al., 2020). The increased difficulty likely stems from more broadly similar stimuli within some sound-classes: the same speech utterances from different speakers of the same gender and multiple human environmental sounds from similar media, both of which were frequently confused. Participants also expressed a moderate amount of difficulty identifying the instrument sounds similar to previous studies suggesting these three categories were perhaps better equated for recognition difficulty. Twenty-seven participants reached the 80% criterion on their first attempt, fourteen participants required a second attempt at screening before reaching criterion, and one participant required a third. Participants were not introduced to the synthesized stimuli prior to the scan, since any identification label given to them would have been arbitrary. However, before the scan participants were told they would infrequently hear some additional synthesized sounds that they did not hear at screening (described as “computer tones and swishes of noise”) and that they should simply listen and pay attention to these like all the other stimuli with no special response required.

At the EEG session, participants were fitted with a cap and electrodes for a 64-channel Biosemi Active-Two system (10-20 system layout) with six additional electrodes applied to record the left and right mastoids as well as left and right horizontal and vertical EOG. Offsets for the scalp sensors were kept within +/− 40 mV. EEG data was recorded at a sampling rate of 512 Hz. During the EEG session, participants watched a nature documentary with the audio muted. Participants were instructed to pay attention to the sound stimuli and to try to limit their movements during the presentation of the sounds. To ensure that participants remained attentive to the sounds they were asked to make a button press following each dog vocalization stimulus.

Stimuli were presented over the course of 4 runs of trials. During a run, participants heard each natural sound token 20 times (80 trials total across runs), each synthesized stimulus 5 times (20 trials total across runs), and the dog vocalization stimulus 70 times. Trials were presented at a jittered rate of one per second (+/− 200 ms). A small number of playback errors resulted in shortened ISIs. Thus, any pairs of trials with less than a 600 ms ISI were removed prior to analysis (0.6% of the data).

#### 4. Data Analysis

The 64 scalp EEG channels were first re-referenced to their common average. The data from each participant was then converted into principal components (retaining 99.9% of the variance) to provide the classifier with a set of orthogonal features (note, essentially the same results were obtained using the raw EEG data instead of principal components). Trials were then epoched around the onset of each stimulus from −200 to 600 ms, and down-sampled to 256 Hz. Edge artifacts from down-sampling were eliminated by removing the extra 100 ms on either side of the −100 to 500 ms epoch. Each EEG channel was then corrected to the average of its 100 ms pre-stimulus baseline period.

Decoding and representational similarity analyses of the natural stimuli were carried out as in Ogg et al., 2020. Each unique pair of items (630 sound token pairs, one half of the representational dissimilarity matrix, or “RDM”) was decoded at each time point throughout the time series at the individual subject level (via CoSMoMVPA; Oosterhof et al., 2016) using a linear discriminant classifier and 90/10 cross validation. The principal components (derived from the EEG) at a given time step were used as feature vectors for training and testing.

Subject-level averages over cross validation folds and specific sections of participants’ neural RDMs were submitted to group level analyses of classification rates for overall decoding as well as individual within and between class comparisons. Group level analyses of decoding accuracy were carried out using non-parametric bootstrapped tests (Maris & Oostenveld, 2007) and threshold free cluster enhancement across time points in the epoch (Smith & Nichols, 2009) using CoSMoMVPA (Oosterhof et al., 2016) to asses when decoding exceeded chance (using permutation sign tests against chance) or when decoding rates differed between comparisons (using bootstrapped t-tests or ANOVA).

Acoustic features of the natural stimuli were extracted and the pair-wise differences among the stimuli for each feature (absolute or Euclidean distances) were organized into acoustic RDMs for correlations against the neural RDMs via representational similarity analysis (Kriegeskorte and Kievit, 2013) using the same methods described by Ogg and colleagues (2020). See Table 1 for a description of each feature along with notes regarding the calculation and interpretation of each. Temporally varying features for the local acoustic analysis were extracted in 5 ms increments. The global acoustic analysis comprised features that characterized each stimulus across its duration (temporal characteristics such log-attack-time and temporal centroid or modulation power spectra), or the median value across windows of the time-varying features. Neural RDMs were correlated with (global or local) acoustic RDMs at the individual subject level using Kendall’s tau correlations. Subject-level correlation statistics were analyzed at the group level via Wilcoxon signed-rank tests against zero, and false discovery rate corrected (*p* < 0.001, Benjamini & Hochberg, 1995) across time points.

In addition to the acoustic RDMs, behavioral dissimilarity ratings for each pair of sounds were collected from a group nineteen participants (who did not participate in the EEG study or Experiment 1). Participants provided pairwise dissimilarity ratings using the same protocol as Ogg and Slevc (2019a). These ratings were averaged into a single behavioral RDM that was analyzed in the same manner as the global acoustic features.

To provide more relevant context for class responses to the synthesized stimuli, an additional classification analysis was run on the natural stimuli to specifically look at three-way, sound-class decoding accuracy (i.e., three-way decoding of super-ordinate sound-class labels, not pair-wise decoding between individual sounds, thus chance classification accuracy = 1/3). This used a similar 90/10 cross validation scheme across trials, but was carried out using the neural responses for all the natural sounds within each fold. This again yielded 10 folds of classification accuracies for each subject at each time point, which were averaged by subject and analyzed using the same group-level tests against chance in CoSMoMVPA.

Synthesized stimuli were analyzed using a modification of the above (three-way) classification approach at the individual subject level. However, instead of using 90/10 cross-validation, the classifier was trained on the neural responses and class labels of all the trials containing natural sounds at a given time point. The classifier was then “tested” on the neural response for each trial containing a synthesized stimulus (at the same time point) and the label output by the classifier was recorded. This elicited a prediction from the classifier about what sound class the neural responses to the synthesized stimuli resembled relative to the responses to the natural stimuli in the training data. This classification scheme was carried out for each synthesized stimulus at each time point for each participant.

The classifier’s predictions for the synthesized stimuli were statistically analyzed using generalized binomial mixed effect regressions similar to Experiment 1 (with a logit link function using ‘lme4’ in R; Bates et al., 2015; statistical significance assessed with ‘lmerTest’; Kuznetsova et al., 2017). These models were fit to the classifier’s trial-level predictions for each synthesized stimulus throughout the EEG timeseries. Again, this involved three one-hot-coded binary outcome variables that indicated whether the classifier made a prediction for each class sound-class (1) or not (0). A separate model was fit for each of the binary sound-class outcome variables and for each of the three manipulation conditions (aside from aperiodicity which interacted with each), for a total of nine models. Again, each of these models was fit only to the subset of trials relevant to each manipulation. To simplify the modeling analysis, every five time points following the onset of significant three-way class decoding was retained (approximately every 20 ms). These models contained predictors for the time point in the epoch (z-normalized), the aperiodicity condition (i.e., the acoustic carrier with 2 contrast coded levels: low aperiodicity/sawtooth and high aperiodicity/noise, with the sawtooth condition as the reference level; corresponding to rows in Figure 1), and a predictor for one of the other manipulations: the spectral centroid, spectral variability or fundamental frequency factors (each with three levels: low, medium, or high, dummy coded with low designated as the intercept; corresponding to columns in Figure 1). The two and three-way interactions among these variables were also included. Note that again, models of the fundamental frequency manipulation did not contain the aperiodicity predictor because these all used the sawtooth wave carrier and there was no noise stimulus for that condition. The random effects structure of all these models contained slopes for time points nested with random intercepts for participants and the slopes and intercepts were constrained to be uncorrelated. Thus, aside from differences in how duration was handled and coded, essentially the same binomial mixed effect models were fit to participants’ behavioral identification responses in Experiment 1 and the classifier’s predictions for the synthesized stimuli in Experiment 2.

### B. Results

Participants in Experiment 2 heard 80 repetitions of 36, 300 ms natural sound tokens evenly divided between three super-ordinate classes: speech, musical instrument and human environmental. Interspersed with these natural sounds was a set of synthesized stimuli designed to parametrically manipulate the acoustic features associated with these sound classes in prior neural and behavioral studies. Findings regarding the natural sound stimuli are reported first. As in Experiment 1, re-examining previous decoding results for the natural sounds contextualizes and validates the approach of manipulating relevant acoustic dimensions via the synthesized stimuli.

#### 1. EEG decoding of the natural sound tokens

Individual pairs of natural sound tokens were first decoded from one another throughout the EEG timeseries at the individual subject level. Decoding rates across pairs of stimuli throughout the epoch are summarized (averaged across participants) in the neural RDMs in Figure 7. Subsections of these RDMs within or between categories (or across the entire RDM to assess overall decoding accuracy) were then averaged for each subject and these subject-level averages were analyzed at the group level. EEG decoding accuracy for this set of sounds followed a similar pattern seen in prior MEG work (Ogg et al., 2020) with a peak around 150 ms (overall peak EEG decoding = 54.1% at 148 ms) followed by a sustained period of lower but above chance accuracy throughout the rest of the epoch (500 ms) past sound offset at 300 ms. This cluster of time points with significant above-chance EEG decoding across all stimuli began at 47 ms (Figure 7B top; assessed via bootstrapped sign-permutation testing and threshold-free cluster enhancement across time points, all *p* < 0.05 corrected).

Decoding pairs of stimuli both within and between sound categories from EEG responses significantly exceeded chance as well, however (as observed in the MEG data) between-class decoding accuracy was higher than within class decoding accuracy within a large cluster of time points beginning at 70 ms and lasting throughout the rest of the epoch (Figure 7B bottom; assessed via non-parametric bootstrapped *t*-tests for between vs. within-class accuracy and threshold-free cluster enhancement across time points, all *p* < 0.05 corrected). Clusters of significant (all *p* < 0.05, corrected) between-class decoding for human-environmental vs. instrument (47 to 500 ms; peak = 57.1% at 148 ms), human-environmental vs. speech (51 to 500 ms; peak = 53.8% at 148 ms) and instrument vs. speech (51 to 500 ms; peak = 54.5% at 145 ms) sounds were observed as well. Human-environmental vs. instrument sound pairs were the most accurately decoded (clusters of significant differences among between class decoding: 121 to 187 ms, 227 to 277 ms, and 348 to 500 ms assessed via non-parametric bootstrapped ANOVA and threshold-free cluster enhancement over time points, *p* < 0.05, corrected). Clusters of significant within-class decoding for speech (largest cluster: 105 to 449 ms; peak = 52.6% at 336 ms), instrument (largest cluster: 78 to 430 ms; peak = 52.3% at 188 ms) and human environmental sound pairs (82 to 500 ms; peak = 51.9% at 184 ms) also exceeded chance (all *p* < 0.05, corrected). Speech decoding was also the most accurate among the within class comparisons, although the peak came later in the epoch (largest cluster of significant differences among within class decoding 289 to 382 ms, *p* < 0.05, corrected).

Three-way decoding for the superordinate class labels associated with each of the stimuli was also significant (i.e., decoding whether the sound on a given trial was speech, music or human-environmental). This is similar to the training dataset and classification procedure used for the analysis of the synthesized stimuli, and thus provides context for how well the classifier might assign these labels to the synthesized sounds. The results of this classification analysis are displayed in Figure 8. The time-course of these results is similar to the pattern of overall decoding (Figure 7B). Superordinate class decoding exceeded chance (chance = 1/3) in a cluster of time points beginning at 51 ms and lasting throughout the epoch (500 ms). Peak three-way class decoding accuracy was 40.3%, or about 7% better than chance, and occurred at 148 ms. Subjects’ level of musical training or behavioral performance at screening was not correlated with decoding accuracy across subjects at any time point (overall, in Figure 7B top, or for any combination of categories in Figure 7C or 7D, or for three-way class decoding in Figure 8, all FDR corrected p > 0.05 across time points for each measure).

**Figure 8.**
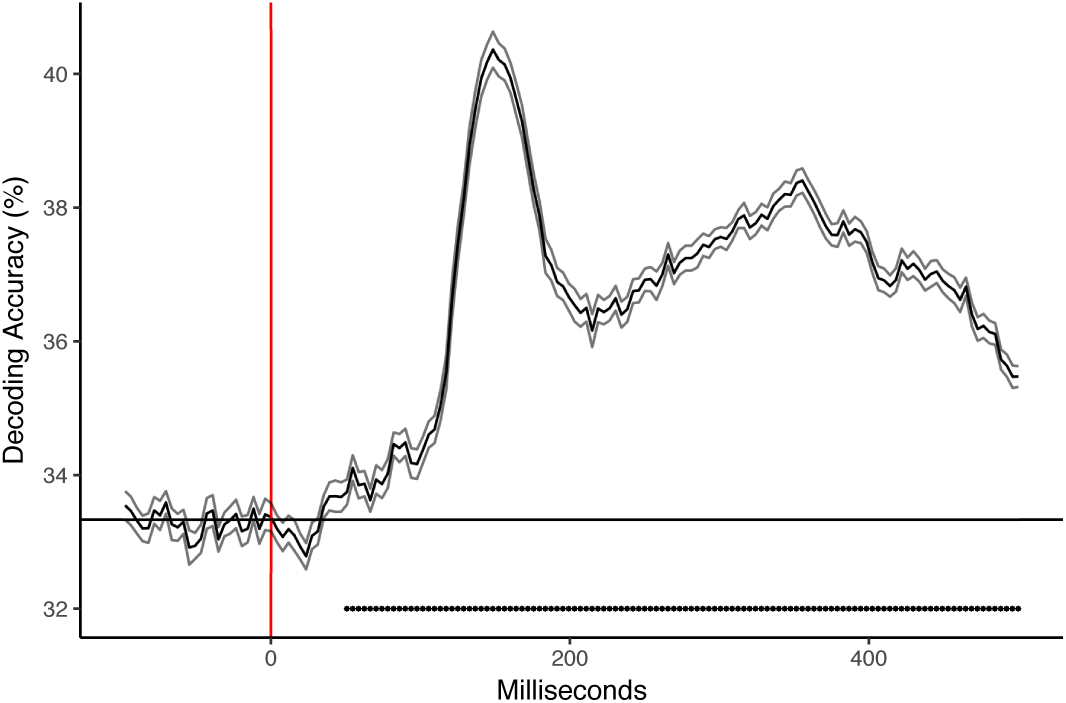
Grand averaged three-way decoding accuracy for super-ordinate sound-class labels of the natural stimuli in Experiment 2 (speech, music or human-environmental; * p < 0.05 corrected v.s. chance = 33.3%). Lines indicate +/– one standard error.

To obtain a high-level view of the time course of the relationship between these decoded neural representations and behavioral impressions of these stimuli, nineteen additional participants provided pairwise dissimilarity ratings of these sound stimuli using the same procedure as Ogg & Slevc (2019a). These ratings were averaged into a behavioral RDM, which was submitted to a representational similarity analysis that correlated this behavioral RDM with participants’ neural RDMs throughout the EEG time series (Figure 9). The largest cluster of time points where the behavioral and neural RDMs were significantly correlated extended from 105 ms to the end of the epoch at 500 ms (FDR corrected *p* < 0.05 across timepoints). Behavioral ratings were maximally correlated (*r_τ_* = 0.14) with the neural RDM at 145 ms.

**Figure 9.**
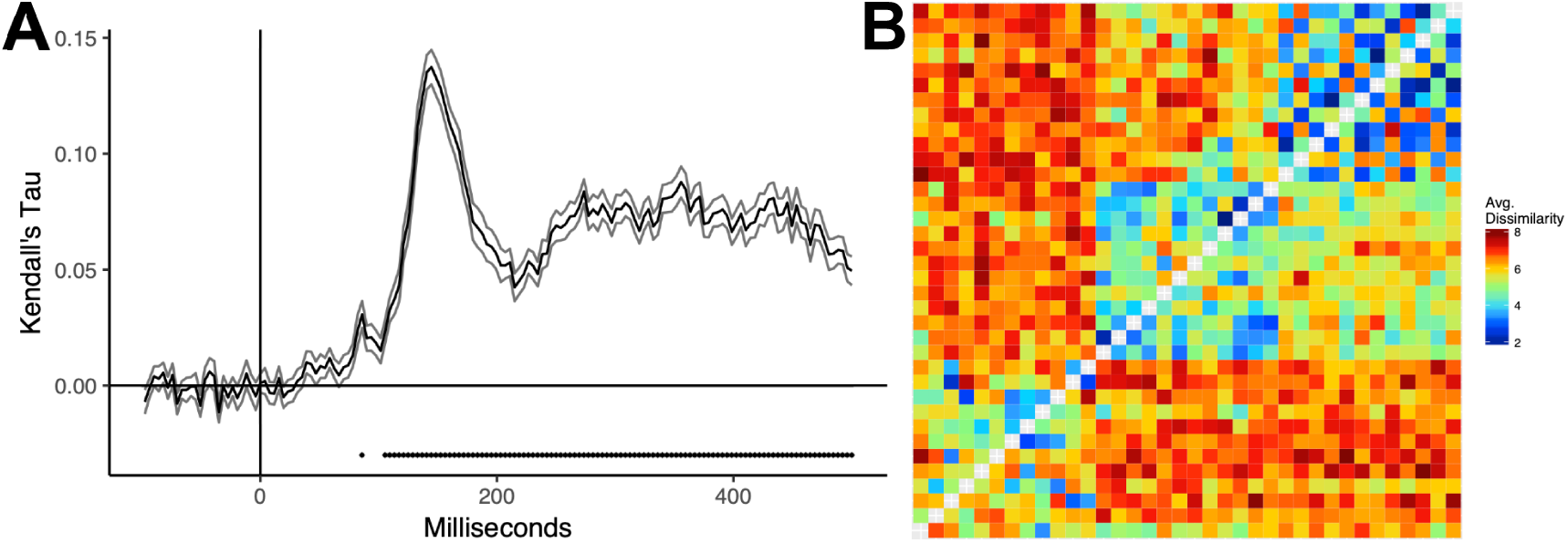
Correlation between EEG decoding accuracy rates (neural RDMs) and behavioral dissimilarity ratings (behavioral RDM) of the natural stimuli throughout the epoch. (A) Mean correlation statistics and standard errors across participants are plotted throughout the neural epoch (* FDR-corrected *p* < 0.001). (B) Behavioral RDM of the pairwise dissimilarity ratings for each stimulus average over ratings obtained from 19 participants using the same protocol as Ogg and Slevc (2019a).

To understand what acoustic features were related to the decoding patterns that emerged throughout the EEG timeseries, the neural RDMs were submitted to representational similarity analyses that correlated these decoding rates with global (aggregated across the whole stimulus duration; Figure 10A) and local (5 ms windows throughout the stimuli; Figure 10B) acoustic RDMs corresponding to features of the stimulus pairs (see Table 1; Methods). In general, the same global acoustic features identified in previous work using MEG (Ogg et al., 2020) correlated with these neural decoding results (all FDR-corrected *p* < 0.001) although the EEG results revealed a more reliable influence of spectral flatness, spectral centroid and ERB Energy (which could be attributable either to the different modalities or stimulus sets). The earliest correlations observed between the global acoustic features and the neural RDMs were for the overall ERB spectral envelope (at 94 ms in the neural epoch, *r_τ_* = 0.03), modulation power spectra (at 105 ms, *r_τ_* = 0.02), and spectral variability (at 113 ms, *r_τ_* = 0.03). The strongest correlations between the neural RDMs and the global acoustic features were for aperiodicity (at 148 ms in the neural epoch, *r^τ^* = 0.12), modulation power spectra (at 148 ms, *r_τ_* = 0.11), and spectral variability (at 145 ms, *r_τ_* = 0.09).

**Figure 10.**
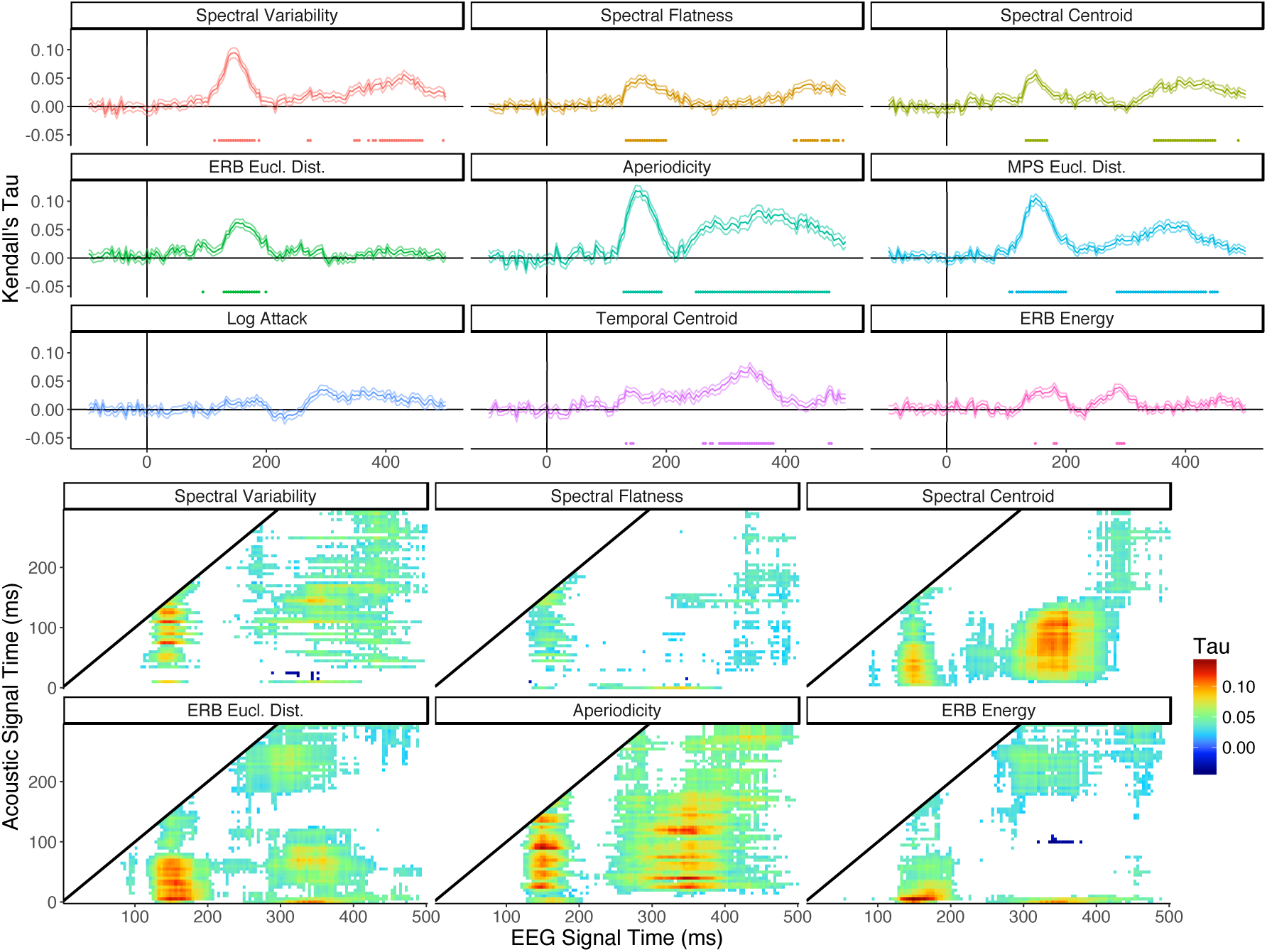
Correlation between EEG decoding accuracy rates (neural RDMs) and acoustic feature differences of the natural stimuli (acoustic RDMs) throughout the epoch plotted as averages of subject-level correlations. (A) Global acoustic analysis results. Mean correlation statistics and standard errors across participants are plotted for each feature throughout the neural epoch (* FDR-corrected *p* < 0.001). (B) Local acoustic analysis results. Mean correlation statistics across participants at the neural and acoustic time points that were significant at the group level (FDR-corrected *p* < 0.001) are shown.

Local acoustic feature analyses revealed significant correlations (all FDR-corrected *p* < 0.001) between some of the earliest time points in the stimuli and the neural decoding patterns (in particular, for ERB Energy: 5 ms in the stimulus correlated with 55 ms in the neural epoch; *r_τ_* = 0.03). The strongest correlations observed in the local acoustic analyses were for ERB Energy (between 5 ms in the stimulus and 148 ms in the neural epoch, *r_τ_* = 0.15), aperiodicity (between 90 ms in the stimulus and 148 ms in the neural epoch, *r_τ_* = 0.14), and the overall spectral envelope (or ERB cochleagram, between 5 ms in the stimulus and 148 ms in the neural epoch, *r_τ_* = 0.13).

Thus, a very similar pattern of results relative to previous work (Ogg et al., 2020) was obtained for decoding this new set of auditory objects and events from EEG responses. These similarities come alongside some notable differences in timing (earlier above chance decoding was achieved in EEG) and accuracy (lower overall decoding rates and correlations with acoustic features in EEG), and the relative influence of some acoustic features. Nonetheless, this broad similarity with prior results in terms of the pattern of decoding accuracy and the most influential acoustic features provides a good grounding for subsequent analyses of the synthesized stimuli, which were designed to parametrically manipulate these same acoustic qualities.

#### 2. Analyses of synthesized sound tokens

Having obtained a predicted pattern of decoding results (both in-terms class-level comparisons and associations with key acoustic features), the approach of manipulating acoustic dimensions via the synthesized stimuli will be informative. Sound-class predictions (i.e., a speech, musical instrument, or human-environmental label output) were elicited from the classifier for each synthesized stimulus and these are summarized in Figure 11. Predictions were obtained after training the classifier on the sound class labels and neural responses for the natural stimuli at a given timepoint and “testing” the classifier on each synthesized stimulus trial at the same time point (again, at the individual subject level and proceeding throughout the EEG timeseries). Although there is no technically correct label for the synthesized stimuli on these “test” trials, the label that the classifier assigns provides an indication of how closely the neural response to each of the acoustic manipulations resembles neural responses to the natural sounds from these categories.

**Figure 11.**
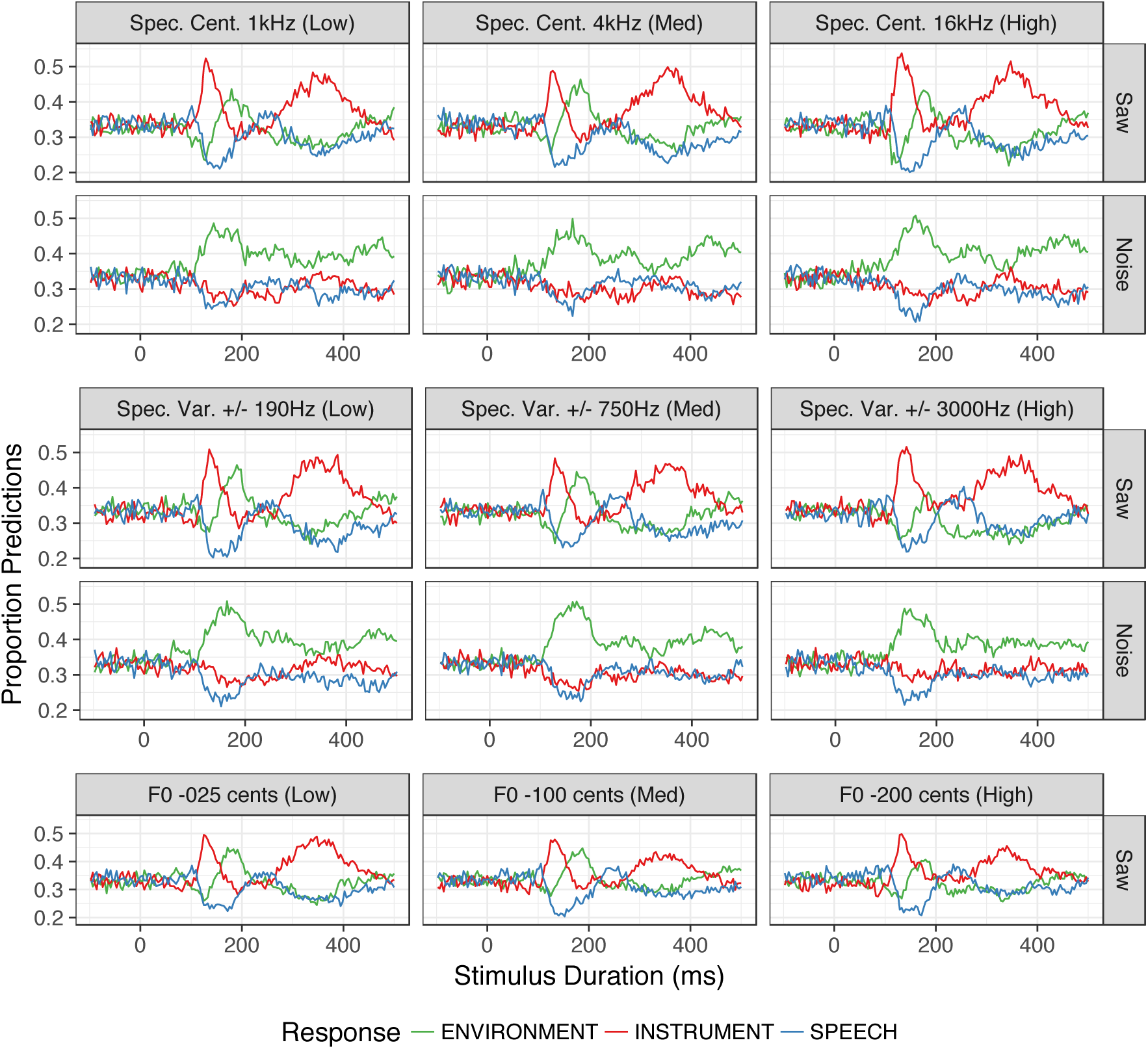
Summary of the classifier’s class predictions for synthesized stimuli throughout the EEG epoch in Experiment 2. Predictions are plotted as a function of the acoustic manipulation conditions (same layout as Figure 1) and averaged over participants and instances of each synthesized stimulus.

Figure 11 demonstrates that as expected, prior to sound onset and early in the epoch the classifier is equivocal in its assignment of class labels: all categories are assigned at similar rates close to chance (1/3), and thus no class-related information is present in the response. Later in the epoch, however, the rates at which the classifier assigns each class label begin to diverge and the resulting pattern is reminiscent of trends observed in the behavioral responses to the synthesized stimuli in Experiment 1 (Figure 6) as well as the pattern of three-way class decoding among the natural sounds (Figure 8). Again, the aperiodicity manipulation appears to exert a large influence on the rates of human-environment (high aperiodicity, noise source) and instrument (low aperiodicity, sawtooth wave source) predictions provided by the classifier. Speech stimuli also appear to be poorly characterized by the synthesized sounds and mostly resulted in speech response rates at or below baseline. Similar to Experiment 1, manipulations of spectral centroid, spectral variability and fundamental frequency appear to exert a more subtle influence on prediction rates.

To examine the influence of these acoustic manipulations on the classifier’s prediction rates across time, a series of generalized binomial mixed effect regressions were fit to the binarized class response variables (i.e., one-hot encoding the class predictions). These predictions were analyzed as a function of the corresponding time point in the epoch (analogous to the duration predictor in Experiment 1) and the aperiodicity condition crossed with the spectral centroid, variability or fundamental frequency manipulations. Note, this analysis only retained time points after three-way class decoding significantly exceeded chance for the natural sounds (51 ms), since this is when sound class information was present in the neural responses.

These models are summarized in Table 6 and generally support the observations above. Across all manipulations, the instrument predictions were associated with low aperiodicity (sawtooth wave carrier), while the human-environmental predictions were associated with high aperiodicity (noise carrier). The fundamental frequency variability manipulations reduced instrument-class prediction rates (especially later in the epoch: negative main effect of medium fundamental frequency variability and negative interactions between fundamental frequency variability and time point) and increased human-environmental-class prediction rates (positive main effect of medium fundamental frequency variability). Greater fundamental frequency variability also increased speech-class predictions but to a lesser degree than for the other categories and only in the presence of the highest fundamental frequency variability (small positive effect of high fundamental frequency variability), and speech prediction rates were still generally lower than baseline. The highest spectral centroid condition with a noise carrier increased human-environmental prediction rates (positive interaction between noise and high spectral centroid). Speech prediction rates were slightly increased for the medium spectral centroid condition with a noise carrier (positive interaction between noise and medium spectral centroid), and were reduced at later points in the epoch given the highest spectral centroid condition (negative interaction between high spectral centroid and time point). Instrument responses were attenuated at later time points for the noise and medium spectral centroid condition (negative three-way interaction). Instrument predictions were generally reduced with spectral variability especially with the noise carrier (negative main effect of medium spectral variability and negative interactions between noise and spectral variability). Greater spectral variability generally increased speech prediction rates (positive main effects), but reduced them at the highest variability when paired with a noise carrier (negative interaction between noise and high spectral variability). High spectral variability generally decreased human-environmental prediction rates (negative main effect for the high spectral variability), but the exception to this was the noise carrier in the high spectral variability condition, which elicited more human-environmental predictions (positive interaction between noise and high spectral variability).

**Table 6.**
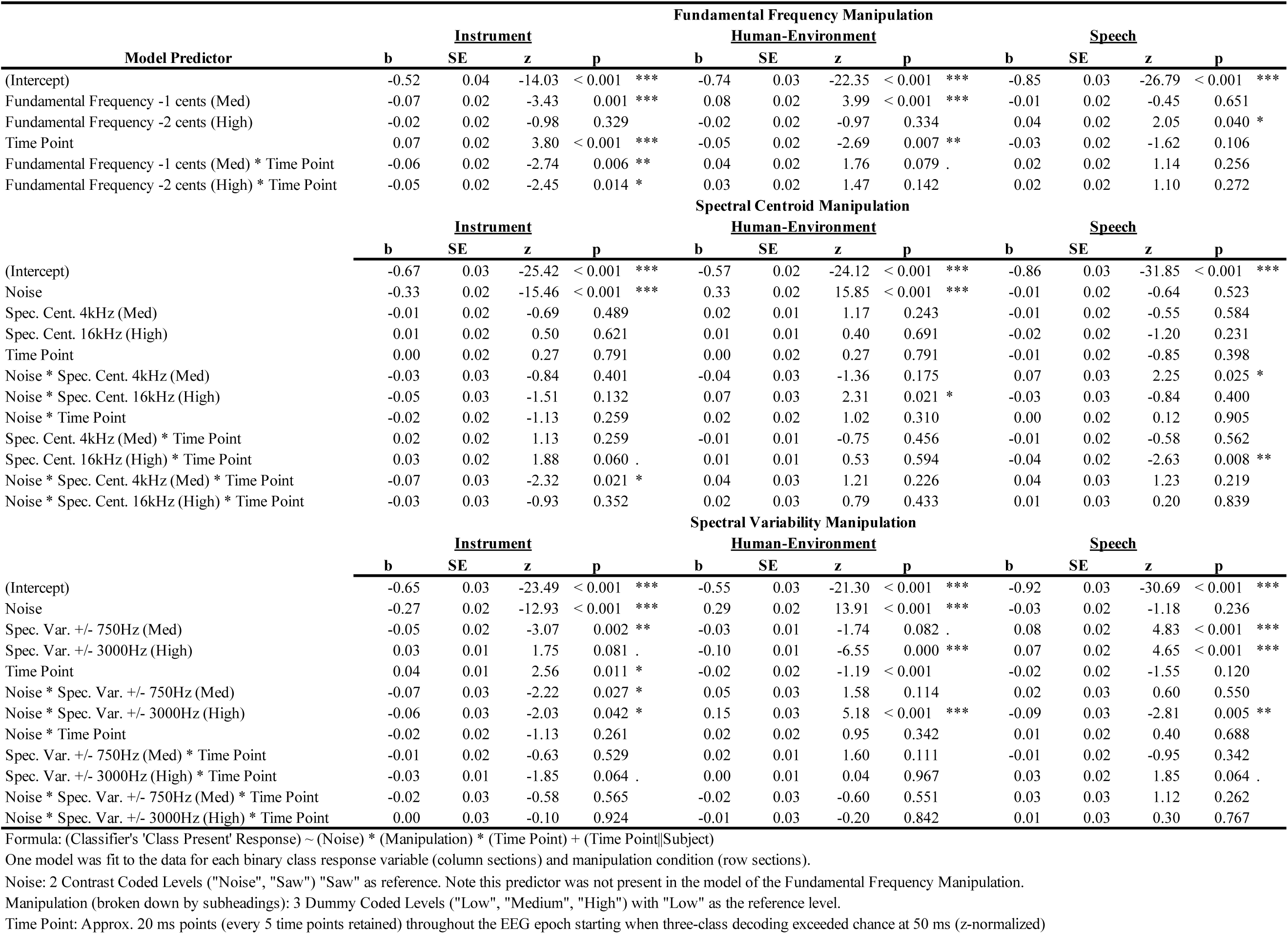
Experiment 2, Models of Predictions from the Pattern Classifier for Synthesized Stimulus Manipulations.

### C. Experiment 2: Discussion

Experiment 2 decoded EEG responses for a set of natural sound tokens as well as a set of synthesized sounds designed to manipulate acoustic features identified in previous work. Results based on the synthesized stimuli converged with many key hypotheses and findings from correlational analyses. Along with Experiment 1 this work underscores the importance of noise cues and to a slightly lesser degree spectral variability, spectral envelope, and fundamental frequency cues in the formation of neural representations of incoming sound stimuli.

Decoding results for the natural stimuli in EEG generally aligned with previous findings using this approach (in MEG: Ogg et al., 2020), which paved the way for tests using synthesized stimuli that directly manipulated aperiodicity, spectral centroid, spectral variability and fundamental frequency cues implicated in earlier work. This approach largely converged with acoustic analyses of responses to the natural stimuli based on representational similarity analysis, particularly for the influence of aperiodicity on the instrument and human-environmental predictions. The fundamental frequency manipulation results also aligned with predictions for the instrument stimuli. However, the influence of spectral centroid was less pronounced in this study compared to Experiment 1. The spectral variability results mostly aligned with predictions for the speech stimuli, and partially aligned with predictions with the human environmental stimuli and instrument stimuli. Generally, the spectral centroid and spectral variability manipulations appeared to exert a moderating influence on the aperiodicity of the carrier (i.e., these effects were often observed as interactions between feature manipulations and aperiodicity). Taken together, these findings further emphasize the importance of harmonic, spectral (and temporal) envelope, spectrotemporal, and fundamental frequency cues on the emerging neural representations of auditory objects and events.

One of the major differences between these EEG decoding results and prior MEG results (Ogg et al., 2020; Lowe et al., 2020) was the observation of lower overall decoding accuracy for the natural stimuli in EEG. This might have subsequently affected the strength of the correlations that were obtainable in the representational similarity analyses, and potentially introduced some noise into the pattern classifier’s predictions for the synthesized stimuli. Reduced EEG decoding accuracy could result from a variety causes: 1) fewer features available for training and testing (i.e., 64 EEG channels v.s. 157 MEG channels), 3) fewer trials for training and testing due to time constraints to accommodate the synthesized stimuli (80 repetitions of each stimulus in the EEG study v.s. 100 repetitions in the MEG study), 3) lower spatial resolution of the EEG signal itself (Baillet, 2017), or 4) from differences among the stimulus sets used in the two studies. Indeed, participants in this EEG study had greater difficulty perceiving differences among these sounds at screening than was the case in previous work (Ogg et al., 2020). This was particularly true for distinguishing speakers of the same gender from one another and for the human-environmental sounds from the same excitation media. Despite these ostensible differences, this EEG decoding study still obtained a strikingly similar overall pattern of results as the MEG study (both in terms of relative class decoding and associations with acoustic features) for a completely new set of stimuli.

Another notable departure of these results from previous M/EEG studies (Charest et al., 2009; Lowe et al., 2020; Lucia et al., 2010; Murray et al., 2006; Ogg et al., 2020), relates to the earlier onset of when super ordinate sound categories could be distinguished from one another. These earlier decoding onsets could be attributed to from stimulus-evoked differences in middle latency response components such as Nb (or TP41; Cacace et al., 1990) and Pb, both of which have a cortical origin (Liegeois-Chauvel et al., 1994; Yvert et al., 2001). Notably these 40-50 ms components may originate in radially oriented sources (Cacace et al., 1990; Scherg & Von Cramon, 1986; Yvert et al., 2001), which are not captured well in MEG (Ahlfors et al., 2010; Baillet, 2017; Yvert et al., 2001). Moreover, these middle latency components have been observed to be sensitive to high frequency, aperiodic stimuli (particularly in EEG), resulting in shorter response latencies (Borgmann et al., 2001) and larger response magnitudes (Reite et al., 1982) for white noise and click stimuli relative to tones. These findings based on synthetic stimuli relate to the between-class EEG decoding results seen here, which found strong early decoding for contrasts between the human-environmental sounds (largely aperiodic stimuli with broadband spectra) and the other two categories. The local acoustic analysis provides further support for this suggestion finding that stimulus differences in ERB power and frequency spectra from the earliest stimulus windows correlating with observed decoding rates.

Additionally, previous univariate findings for EEG responses to diverse natural stimuli (Charest et al., 2009; Lucia et al., 2010; Murray et al., 2006), found differences between vocalizations and man made sounds (similar to the human environment vs. speech decoding comparison in Experiment 2) emerging as early as 70 ms, similar to what we see in these EEG decoding results. Closer inspection of these previous univariate findings suggests that even earlier effects might be observed in the time region around approximately 40 or 50 ms (Figures 2 and 3 in Charest et al., 2009; Figure 8 in Lucia et al., 2010; Figure 2 in Murray et al., 2006), but that perhaps these did not survive multiple comparison corrections. Multivariate pattern analyses, like those used in Experiment 2, are often seen as more sensitive when detecting subtle differences between classes than univariate analyses of overall magnitude or latency (Grootswagers et al., 2017; Haynes & Rees, 2006; Tong & Pratte, 2012). This is because pattern analyses leverage the whole pattern of activation and deactivation across sensors, rather than looking at each sensor individually (Charest et al., 2009) or the aggregated response across sensors (Lucia et al., 2010; Murray et al., 2006).

## IV. GENERAL DISCUSSION

Objects and events in the world create sound waves that the auditory system rapidly transforms into representations of the precipitating phenomena. This process allows listeners to identify and react to stimuli in their environment. The two studies in this report attempted to clarify how this recognition process unfolds for the diverse sound stimuli human listeners are sensitive to. Specifically, this work aimed to identify what qualities of the acoustic signal influence the formation of auditory object and event representations early in perception. The results expand on prior work and further implicated cues related to aperiodicity, spectral (and less consistently, temporal) envelopes, spectral variability and fundamental frequency variability as acoustic qualities that influence emerging object representations within the first hundreds of milliseconds after sound onset.

A series of acoustic manipulations were implemented by synthesizing simple, controlled stimuli that changed along key dimensions indicated by prior results. These stimuli specifically targeted superordinate categories of speech, music and human-environmental sounds. Overall, this approach supported previous correlational results in both a duration-gated classification task (Experiment 1) and an EEG decoding study (Experiment 2), finding an overarching influence of aperiodicity on sound-class response rates for instrument and human-environmental sounds. The other acoustic manipulations (for spectral envelope, spectrotemporal variability and fundamental frequency variability) also produced statistically significant changes in classification responses but these effects were moderate relative to the aperiodicity manipulation. Notably, the synthesis approach did not precipitate a large number of speech responses, suggesting that the speech class was not well characterized by these acoustically simplified sounds. Additionally, analyses of the natural stimuli in Experiments 1 and 2 largely aligned the previous findings both in terms of class discriminability and the influence of different acoustic dimensions (Gygi et al., 2007; Huang & Elhilali, 2017; Ogg et al., 2017; Ogg & Slevc, 2019a; Ogg et al., 2020).

The psychophysical (Moore, 2012) and neural (Näätänen & Picton, 1987; Roberts et al., 2000) correlates of these cues have been examined for decades, and they have also been hypothesized to play a role in auditory object and event perception across sound classes (Handel, 1995). However, this notion has not been thoroughly examined empirically among a balanced and controlled set of sounds that sample the diverse, natural stimuli humans are sensitive to. Nor has the influence of these features been thoroughly examined during the early time periods when acoustic stimuli and their corresponding perceptual representations are developing following sound onset. Thus, these findings provide new insight into how the auditory system accumulates acoustic information to furnish object and event representations just as they begin to take shape. Taken with estimates that an approximately 25 millisecond lag exists between the auditory periphery and cortex (Liegeois-Chauvel et al., 1994; Raij et al., 2010), reliable behavioral classification results given 25-ms of duration in Experiment 1 and the onset of above-chance EEG decoding around 50-ms in Experiment 2 suggests an approximately 25-ms integration window might exist post stimulus on onset for establishing representations of incoming acoustic stimuli. The behavioral and neural RDM correlations (Figure 9) in Experiment 2 further suggest that by 100-ms to 150-ms (thus, potentially based on the first 75-ms to 125-ms of stimulus exposure) the neural response captures the bulk of a listener’s high-level percept of a given sound. This is because the behavioral portion of this correlation was based on dissimilarity ratings made for the full 300-ms stimuli. These findings also begin to outline a feature space that encapsulates these behaviorally relevant classes of speech, musical instrument and everyday object sounds. These results suggest that that space is outlined at least by dimensions related to aperiodicity, spectral envelope and spectral variability (with additional dimensions that likely capture temporal envelope, fundamental frequency qualities, and speech related cues described in more detail below).

The temporal evolution of these perceptual and acoustic dynamics help contextualize later, potentially class-specific, neural processes that might be cued or engaged by an early sensitivity to certain acoustic features. A large part of our current understanding of how the human brain processes the acoustic features of natural sounds in service of auditory object and event recognition comes from relatively slow responses observed in fMRI. Although the issue of neural overlap between sound categories is still a matter of active debate (particularly for music and language, see Ogg & Slevc, 2019b, Peretz et al., 2015 for reviews), fMRI studies generally find class-dependent response patterns in various parts of temporal cortex (Angulo-Perkins et al., 2014; Leaver & Rauschecker, 2010; Lee et al., 2015; Norman-Haignere et al., 2015; Ogg et al., 2019; Rogalsky et al., 2011; Staeren et al., 2009). This aligns with the high between-class decoding rates and robust between-class behavioral responses observed in Experiments 1 and 2. While some studies suggest certain class-specific responses that are independent of the acoustic features of the stimuli (Leaver & Rauschecker, 2010; Norman-Haignere et al., 2015), many of the features implicated in the Experiments 1 and 2 are also found to influence different areas of cortex in fMRI during natural sound perception (Giordano et al., 2012; Lewis et al., 2009; Lewis et al., 2012; Norman-Haignere et al., 2015). Some go further to suggest that these acoustic feature differences among stimuli can account for observed class-specific responses (Giordano et al., 2012; Giordano et al., 2014; but see Leaver & Rauschecker, 2010; Norman-Haignere et al., 2015). In any case, this report provides additional insight into *when* the auditory system processes different kinds of acoustic information from natural sounds after onset, which may inform how this auditory pathway functions and routes information.

The results of Experiments 1 and 2 raise interesting questions with respect to speech. On the one hand, these stimuli were the most robustly represented among the natural sounds: the speech class that was most accurately discriminated behaviorally, and the sounds within this class were most accurately decoded from one another based on early neural responses. On the other hand, speech appeared to be the least well characterized class of sounds with respect to the synthetic acoustic manipulations. In both Experiments 1 and 2, a notable result for the synthetic tokens was the lack of speech responses (either from participants or the pattern classifier) relative to the other sound categories. This occurred despite having directly manipulated features that previous acoustic analyses (Gygi et al., 2007; Huang & Elhilali, 2017; Ogg et al., 2017; Ogg & Slevc, 2019a; Ogg et al., 2020) implicated as being important for recognizing speech: spectrotemporal variability (particularly for the spectral centroid), and fundamental frequency variability.

How might these findings be reconciled and what can they tell us about speech? One likely possibility is that a feature was missing from the synthesized stimuli that specifically characterizes speech (Johnson, 2005; Johnson, 2011). This missing cue (e.g., formant structure or vocal tract cues) may not have been uncovered by the high-level acoustic analyses designed to characterize a broad variety of sounds (Ogg et al., 2017; Ogg & Slevc 2019a; Ogg et al., 2020). Follow up work might unpack these issues further with studies or manipulations designed specifically with formants in mind.

Another critical aspect of natural speech is that the acoustic cues examined here are rapidly and even *simultaneously* altered within a speech signal. This includes differences in noisiness between voiced and unvoiced segments of speech (Bachorowski and Owren, 1999), emphases on different parts of the frequency spectrum among fricatives or vowels (Johnson, 2011; Ladefoged & Johnson, 2015), transitions or coarticulation between consonants and vowels (Holt & Lotto, 2010), and prosodic cues (Gay, 1978; Grandjean et al., 2006; Heffner & Slevc, 2015). Each of these aspects of speech parallels the aperiodicity, spectral centroid, spectrotemporal and fundamental frequency variability dimensions implicated by the acoustic analyses and altered among the synthesized sounds in these experiments. Indeed, each of these acoustic dimensions might be a parallel channel by which speech transmits information for the listener. Furthermore, this variety of cues (or of information channels) might contribute to the robustness of speech in the presence of acoustic interference. However, this parallelism disagrees with the synthesized stimuli and feature manipulation approach taken in Experiments 1 and 2 that focused instead on isolating individual cues while controlling others. An exciting possibility, given earlier work demonstrating how well attuned speech is to a mammalian auditory system (Smith & Lewicki, 2005), is that our sensitivity to the features explored in Experiments 1 and 2 arose in our ancestors (Town & Bizley, 2013) because these features were useful for auditory object and event recognition, which helped their survival by avoiding threats in the environment. Speech, in turn, may have developed to exploit these acoustic sensitivities to maximize the amount of information that could be transmitted between interlocutors.

## V. CONCLUSION

The ability to recognize objects in the environment is an exceptional feat for our perceptual and cognitive systems. The non-linear transformation of physical stimulation into mental constructs allows us to efficiently navigate a world that is populated by coherent items and events rather than disparate phenomena. It has even been suggested that object recognition is a central problem that our sensory systems have evolved to efficiently accomplish (DiCarlo, Zoccolan & Rust, 2012; Simoncelli & Olshausen, 2001; Yamins et al., 2014). Indeed, the speed and ease with which the world coheres around us (very often prior to our needing to devote any additional mental effort to identifying specific stimuli) is a testament to how impressive this faculty is (DiCarlo & Cox, 2007).

Within just tens of milliseconds, the auditory system accumulates information from acoustic stimuli in the world and begins to mold representations of objects and events. This work has demonstrated that core acoustic cues such as noisiness, spectral (and temporal) envelopes and acoustic change over time (both across parts of the frequency spectrum and for fundamental frequency) guide the auditory system in this process. This set of features subsumes the plethora of stimuli that are important for humans in to a coherent acoustic space.

Casual human listeners might also have some intuitive familiarity with manipulating and parsing this set of cues, given the possibility that speech developed to exploit these diverse acoustic qualities (Johnson, 2011; Mesgarani et al., 2014; Moore, 2012; Smith & Lewicki, 2006; Tang et al., 2017). That is, perhaps speech’s privileged status in human perception owes (at least in part) to its optimal use of the mammalian auditory system’s acoustic sensitivities that developed in our ancestors to support the perception of non-speech objects and events in the world that was critical to their survival. Accordingly, perhaps one reason the synthesized acoustic manipulations of each of these cues did not appear to robustly represent speech is that it exploits a more complex and simultaneous combination acoustic cues that fluctuate from moment to moment. This complexity would not have been captured by the (intentionally) simplified acoustic manipulations employed among the synthesized stimuli here. Instead, these synthesized manipulations are better matched to perhaps more homogenous stimuli that occupy opposing endpoints of our acoustic sensitivities: human-environment and instrument sounds. Indeed, even without invoking potentially speech specific perceptual or neural processes, perhaps one special quality of speech is how it effectively utilizes these acoustic cues as channels to convey information. It will be exciting to further explore this more acoustically unified world sound objects and auditory processes.

## ACKNOWLEDGMENTS

Thanks to Dr. Robert Slevc for feedback and guidance in this work. ChatGPT (GPT-4o, OpenAI) was used for some light copy-editing in the course of preparing this report.

## TEXTUAL FOOTNOTES

1 Note, various iterations of an additional, mixed-aperiodicity condition combining the noise and tone carrier waves was also presented during these experiments. However, results showed that these mixed stimuli were all simply treated (i.e., classified by participants) the same as the noise carrier (observed very similar results to that condition). Thus, for parsimony in reporting these results, and to maximize the power and sample size of these studies, the mixed-aperiodicity trials were removed prior to analysis and participants were pooled across these iterations into one larger sample.

2 This two-stage approach was adopted for three reasons. First, this is a computationally efficient search of the space of possible models in an all subsets regression with interactions among predictors. An exhaustive search of the models containing all combinations of acoustic feature predictors and their interactions with duration involves many thousands of mixed-effects models and many days of computation time. Second, the all-subsets regression algorithm is already constrained so as to only include interactions if both predictors in the interaction are included in the model as predictors (main effects) on their own. Thus, any top ranked model containing interactions between a given feature and duration must perform as well or better than models containing just the top ranked features without the interaction (despite the penalty imposed by the BIC ranking criteria for including additional features). And finally, previous use of an all-subsets regression analysis to analyze results from a gating paradigm (Ogg et al., 2017) found that interactions with duration mainly served to modulate the influence of features that were already strong predictors before the interaction. To confirm the validity of this approach, the data from Ogg et al. (2017) were re-run using the two stage all-subsets regression described here. This obtained the same results as when the set of models and interactions was fully searched by the algorithm.

3 The only significant effects of musical training that were observed were for a lower rate of musical instrument responses for more musically trained participants among the spectral variability stimuli at later durations (interaction between musicianship and duration: *b* = –0.27, *z* = –2.49, *p* = 0.01) and for more speech responses for the noise stimuli with a medium spectral centroid among more musically trained participants (three-way interaction between noise stimulus, medium spectral centroid manipulation and musical training: *b* = 1.23, *z* = 2.02, *p* = 0.04).

